# Digging into the general dynamic model: Island ontogeny predicts shifting micro-evolutionary processes in a Mascarene flowering plant radiation

**DOI:** 10.1101/2025.01.30.635705

**Authors:** Brock Mashburn, Alexander G. Linan, Timothée Le Péchon, Jean Claude Sevathian, Kenneth M. Olsen, Christine E. Edwards

## Abstract

Island studies have been integral in the development of process-oriented biodiversity models such as the general dynamic model (GDM) of oceanic island biogeography. While empirical tests of the GDM primarily come from phylogeographic studies, tests incorporating comprehensive population-level sampling of island radiations are rare. In this study, we elucidate the evolutionary processes driving the diversification of *Hibiscus* section *Lilibiscus* in the Mascarene archipelago using population-level sampling and 2bRAD sequencing. Our goals were to: 1) assess species relationships and resolve taxonomic issues; 2) test patterns of intra- and interspecific genetic differentiation under a model of shifting speciation processes as islands age; and 3) utilize demographic modeling to infer the relative divergence times of populations in the archipelago. We found the Mascarene radiation of sect. *Lilibiscus* to be monophyletic and confirmed the presence of six morphologically distinct species. Species richness and phylogenetic relationships supported the expectations of the GDM. Namely, we found that morphologically similar populations of the same species on the intermediate-aged island, Mauritius, diverged earlier and showed greater divergence than morphologically disparate species on the youngest Mascarene island, Réunion. These patterns are consistent with the hypothesis that ecological selection may be affecting speciation on the youngest island, drift may be the most important force on the middle-aged island. Although not tested here, given that the oldest island is in a late stage of subsidence, extinction may be the most important evolution force occurring on the oldest island, consistent with the expectation that evolutionary processes occurring across each of these islands may be influenced by ontogenetic stage. We also found evidence that some species in the group may be naturally rare, suggesting that the rarity of species may not only be from recent habitat loss. Our study is the first to use population-level data to test complex predictions of the GDM and demonstrate that the most important evolutionary processes shift depending on island ontogeny.

---

Oceanic islands have long served as ideal settings for biologists to study evolutionary patterns and processes. Studies of island radiations have produced processes-oriented models designed to explain patterns of species richness and lineage evolution in island systems. Most famously, the island biogeography theory (IBT) (MacArthur & Wilson 1963, 1967) incorporates the evolutionary processes of dispersal and extinction into predictions of an island’s carrying capacity. Another example is the progression rule (Funk & Wagner 1995), which integrates dispersal with island hot-spot theory to predict a concordance between island age and lineage age in an archipelago, such that older lineages are expected to be found on older islands and younger lineages to be found on younger islands (Borregaard et al. 2017; Cowie & Holland 2006).

The general dynamic model (GDM) of oceanic island biogeography builds on the IBT and the progression rule by incorporating three evolutionary processes — speciation, immigration, and extinction — with geological changes across an island’s life cycle (Whittaker et al. 2008). One of the more commonly tested predictions of the GDM is that of a hump-shaped relationship between species diversity and island age, where middle-aged islands exhibit greater species richness than young or old islands (Prediction 1 in Whittaker et al. 2008). This pattern is assumed to be driven by the interplay between speciation, which is strongest when an island is young and uplifting, and extinction, which predominates as an island ages and subsides. The GDM also makes more complex predictions concerning the evolutionary processes driving diversification at different stages of island ontogeny. However, genomic studies of island radiations testing these process-oriented predictions are rare (Cerca et al. 2023).

According to the GDM, islands exhibit their greatest elevational range, and therefore the highest diversity of ecosystems, soon after uplift (Borregaard et al. 2017; Whittaker & Fernández-Palacios 2007). Thus, in younger islands, ecological speciation is predicted to be more prevalent due to stronger selective pressures across environmental gradients (Prediction 10 in Whittaker et al. 2008). If true, such strong environmental selection would likely lead to quickly-diverging, morphologically distinct species. In contrast, middle-aged islands experiencing erosion and subsidence will exhibit less environmental variation compared to younger islands but may simultaneously display greater topographic complexity as mountains become separated by valleys (Fernández-Palacios et al. 2014), resulting in montane “islands within islands.” If environmental conditions are relatively stable, populations occurring in these “islands within islands” are expected to experience allopatric divergence via drift if they are unable to exchange genes (inferred from Prediction 10 in Whittaker et al. 2008). Moreover, a lack of differential selection pressures among such intraspecific populations would be expected to limit morphological divergence between them, even if these intraspecific populations have been separated for a longer period of time than entire species on younger islands. Finally, old islands in the final stages before submergence are eroded, reducing their geographic size and topographic and environmental diversity. Here, speciation is expected to be rare, and the process of extinction may be the most important evolutionary force affecting species. As relatives that once coexisted on the island are lost through selective or stochastic processes, the remaining species would exhibit greater lineage divergence (longer branch lengths) in relation to each other and to species on the younger islands (inferred from Predictions 5, 7 & 9 in Whittaker et al. 2008).

If the predictions outlined above are true, we can expect different population genetic patterns on islands of different ages within an island chain. For example, species on the younger island should: 1) be the most recently diverged species in a radiation; 2) show less genetic divergence from other species on the same island compared to species pairs occurring on older islands; and 3) be morphologically and environmentally differentiated due to stronger selective forces occurring on younger islands. On a middle-aged island, species should: 1) show intermediate divergence times relative to species on younger and older islands; 2) show greater intraspecific genetic divergence among populations than species on younger islands; and most importantly, 3) exhibit intraspecific populations that are highly diverged genetically without significant morphological or ecological differentiation, due to erosion of the island resulting in the formation of barriers to gene flow, resulting in strong genetic drift. Finally, on the oldest island, species should: 1) occur on long branch lengths and show the earliest divergence times in the entire radiation, in accordance with the progression rule, and 2) exhibit the greatest measures of intraspecific genetic differentiation compared to all other species.

The radiation of *Hibiscus* section *Lilibiscus* in the Mascarene archipelago is an ideal group in which to investigate these predictions. The Mascarene islands are perfect for testing the GDM because of their simplicity in age, size, and geographic distribution. The archipelago consists of three volcanic islands (Fig. 2.1), each in a distinct stage of ontogeny: an old island at a late stage of subsidence (Rodrigues), a middle-aged island experiencing erosion and subsidence (Mauritius), and a young island that is still volcanically active and under formation (Réunion).

Hence, the three islands together exhibit the precise combination of characteristics necessary to test the predictions of the GDM in a tractable way. Recent phylogenetic dating of a global *Hibiscus* phylogeny (Hanes et al. 2024) placed the stem age of sect. *Lilibiscus* at ∼14.66 mya (Maggie Hanes, pers. comm.). This age would make sect. *Lilibiscus* older than the oldest Mascarene island, Rodrigues (∼7.5–11 mya), suggesting that the group arrived in the Mascarenes when Rodrigues was young and diversified as the other islands emerged, a fundamental assumption of the GDM (Duncan 2009; Thébaud et al. 2009).

Another fundamental assumption of the GDM is that a radiation is the result of a single colonization event, and that the taxonomy of a group is well supported. The taxonomy of *Hibiscus* sect. *Lilibiscus* was recently revised based on morphological observations, elevating the number of the species in the group from four to six (Mashburn et al. 2024). Following Mashburn et al. 2024, the Mascarene radiation of sect. *Lilibiscus* consists of six morphologically distinct species distributed among the three islands (Mashburn et al. 2024). As currently circumscribed, the group displays the hump-shaped relationship between island age and species diversity as expected under the GDM, with one species occurring on the oldest island, three on the middle-aged island, and two on the youngest island. However, the group has yet to be analyzed using a phylogenetic approach, such that it is unknown whether genomic data will support the monophyly of Mascarene Lilibiscus and the circumscription of six species rather than four.

Although many studies consider such hump-shaped species diversity pattern to be sufficient evidence that a group evolved according to the processes underlying in the GDM, population occurrence patterns in *Hibiscus* sect. *Lilibiscus* also correspond to the predictions of the GDM. Specifically, two of the species endemic to Mauritius, the middle-aged island, exhibit disjunct populations on ‘islands within islands’ that do not show noticeable morphological or ecological variation. In contrast, the two species endemic to the youngest of island of Réunion are strongly differentiated both morphologically and environmentally. However, it is unclear whether patterns of genetic divergence will also support the expectations of the GDM. Fortunately, the presence of only six species in the Mascarene radiation of sect. *Lilibiscus* permits focused, population-level sampling for population genomic and phylogenomic analyses, which allows for a test of the finer process-oriented predictions of the GDM at inter- and intra-specific scales. The group is therefore poised as a perfect test system for the shifting-process predictions of the GDM.

Our overall objective in this study was to understand the evolutionary history of the Mascarene *Hibiscus* sect. *Lilibiscus* in light of the shifting-speciation process prediction of the GDM. In this study, we performed comprehensive population-level sampling of Hibiscus sect. Lilibiscus on all three Mascarene islands, then generated genome-wide sequence data using a 2bRAD-seq approach (Wang et al. 2012). We reconstructed the phylogenetic relationships of *Hibiscus* sect. *Lilibiscus* in the Mascarenes, analyzed patterns of genetic divergence within and among species, and conducted demographic modeling to understand the relative divergence times of populations and species in the archipelago. We used these analyses to: 1) to test the group’s monophyly and current morphology-based taxonomic hypotheses; 2) test whether patterns of intra- and interspecific genetic differentiation correspond to those expected under a model of shifting speciation processes as islands age; and 3) to determine whether the divergences of morphologically cohesive intraspecific populations in the intermediate-aged island, Mauritius, are more ancient than the divergence between the morphologically disparate species in the youngest island, Réunion, a key component of the predictions outlined above.

## Materials & Methods

### Study system

Located in the Indian Ocean ca. 700–1500 km east of Madagascar (Fig. 2.1), the Mascarene archipelago originates from the southwest to northeast migration of the Central Indian Ridge over the stationary Réunion hotspot (Frey 2009). Rodrigues is the oldest island, forming 7.5–11 mya; the island is in a late stage of subsidence (396 m peak elevation) and is therefore quite small (109 km^2^) with low habitat heterogeneity (Duncan 2009; Middleton & Burney 2013).

Mauritius emerged ∼6.8–7.8 mya, with subsequent periods of limited volcanic activity ending ∼0.1–1.0 mya (Duncan 2009). As a middle-aged island, Mauritius is much larger than Rodrigues (2040 km^2^) and is in a middle stage of subsidence. A ring of small (828 m peak elevation), disjunct mountain ranges are the erosional remnants of the oldest era of volcanic activity and the surrounding valleys resulted from more recent lava flows (McDougall & Chamalaun 1969).

Réunion formed ∼2 mya and continues to experience volcanic activity (Duncan 2009). Réunion is also the largest island of the three islands (2512 km^2^) and has a significantly taller mountain range (3070 m peak elevation) that runs NW to SE through the center of the island. This mountain range creates a much stronger rain shadow effect on Réunion than on the other islands, with very wet (up to 12 m annual rainfall) regions to the east and much drier (∼500 mm annual rainfall) conditions to the west (Thébaud et al. 2009). As a result, Réunion contains a much greater diversity of habitat and forest types than Mauritius or Rodrigues (Cheke & Hume 2008).

In the Mascarenes, following the taxonomy of Mashburn et al. (2024), each of the six *Hibiscus* sect. *Lilibiscus* species is endemic to a single island: *H. liliiflorus* in Rodrigues; *H. dargentii, H. fragilis,* and *H. genevei* in Mauritius; and *H. boryanus* and *H. igneus* in Réunion (Table 3.1, Fig. 3.1). All six species are of conservation concern, and four species — *H. liliiflorus*, *H. dargentii*, *H. fragilis*, and *H. genevei* — have been reduced to two or fewer wild populations (Table 3.1). In Rodrigues, the last two remaining wild individuals of *H. liliiflorus* are found on basaltic soils that cover most of the island, suggesting that the species could have been once been widespread there (Mashburn et al. 2023). Mauritius was relatively well-explored before and during the large-scale deforestation that occurred in the 19th century (Mashburn et al. 2024). Therefore, the absence of collections of *H. dargentii* and *H. genevei* outside of the few extant populations suggests the possibility that these species may not have been widespread or common. The same could be said for *H. fragilis*, which, though now reduced to two populations, was only ever collected from six mountains on the island. In Réunion, *H. igneus* and *H. boryanus* partition the island into two main habitats: *H. igneus* is found in the seasonally dry forests on the leeward side of the island, and *H. boryanus* is found in humid forests on the windward side. Each species is somewhat common within healthy remnant forests of the corresponding habitat type, and it therefore reasonable to infer that each species was once common in similar lower elevation forests that have mostly been lost through deforestation (Albert et al. 2021; Strasberg et al. 2005).

**Table 3.1.**
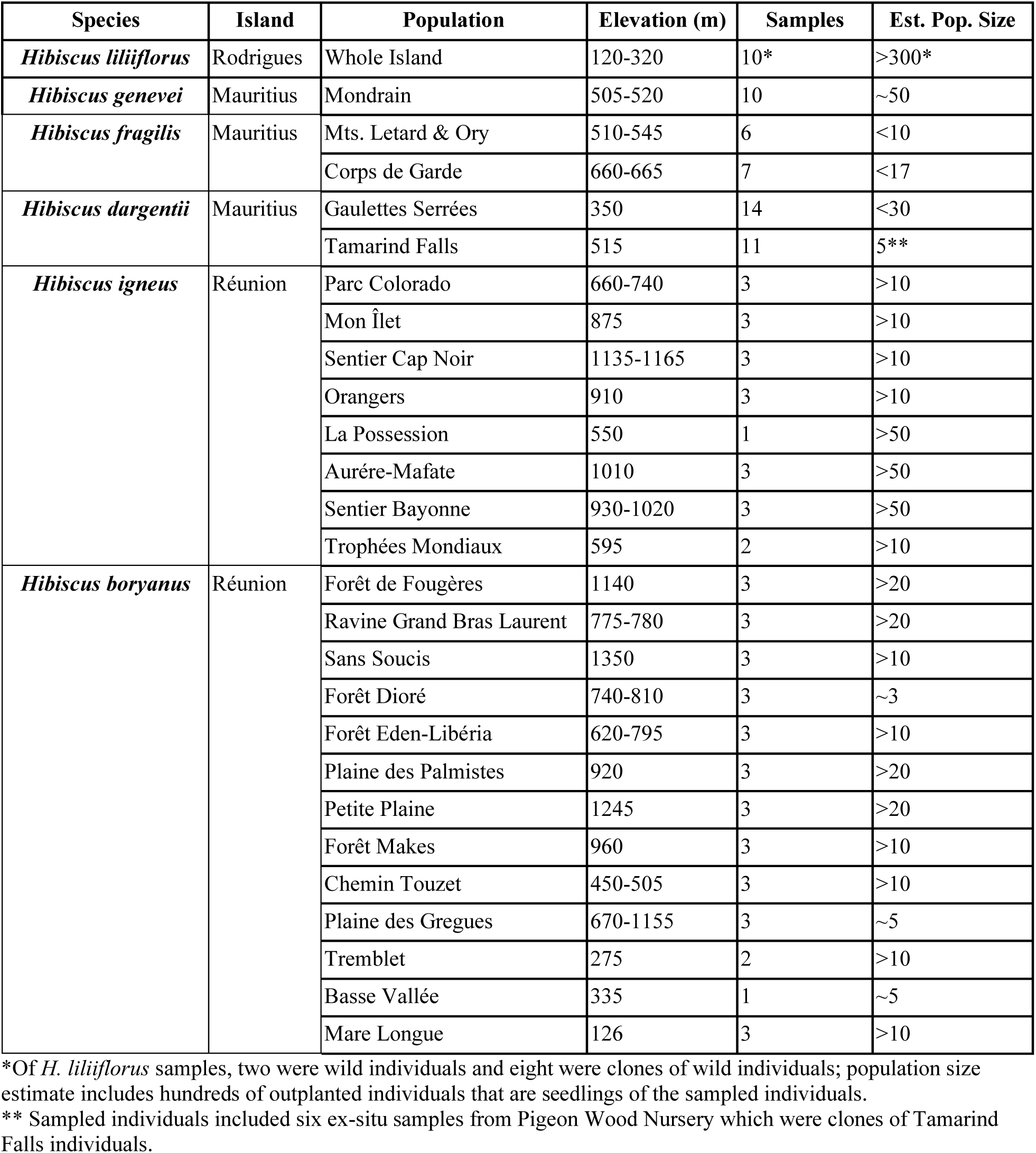
Population information for Mascarene *Hibiscus* sect. *Lilibiscus* collections used in this study. Species are ordered according to island age and phylogenetic position. Populations are ordered from north to south for each species (see Fig. S3.1). Elevation information is based on collections made and included in this study.

**Figure 3.1.**
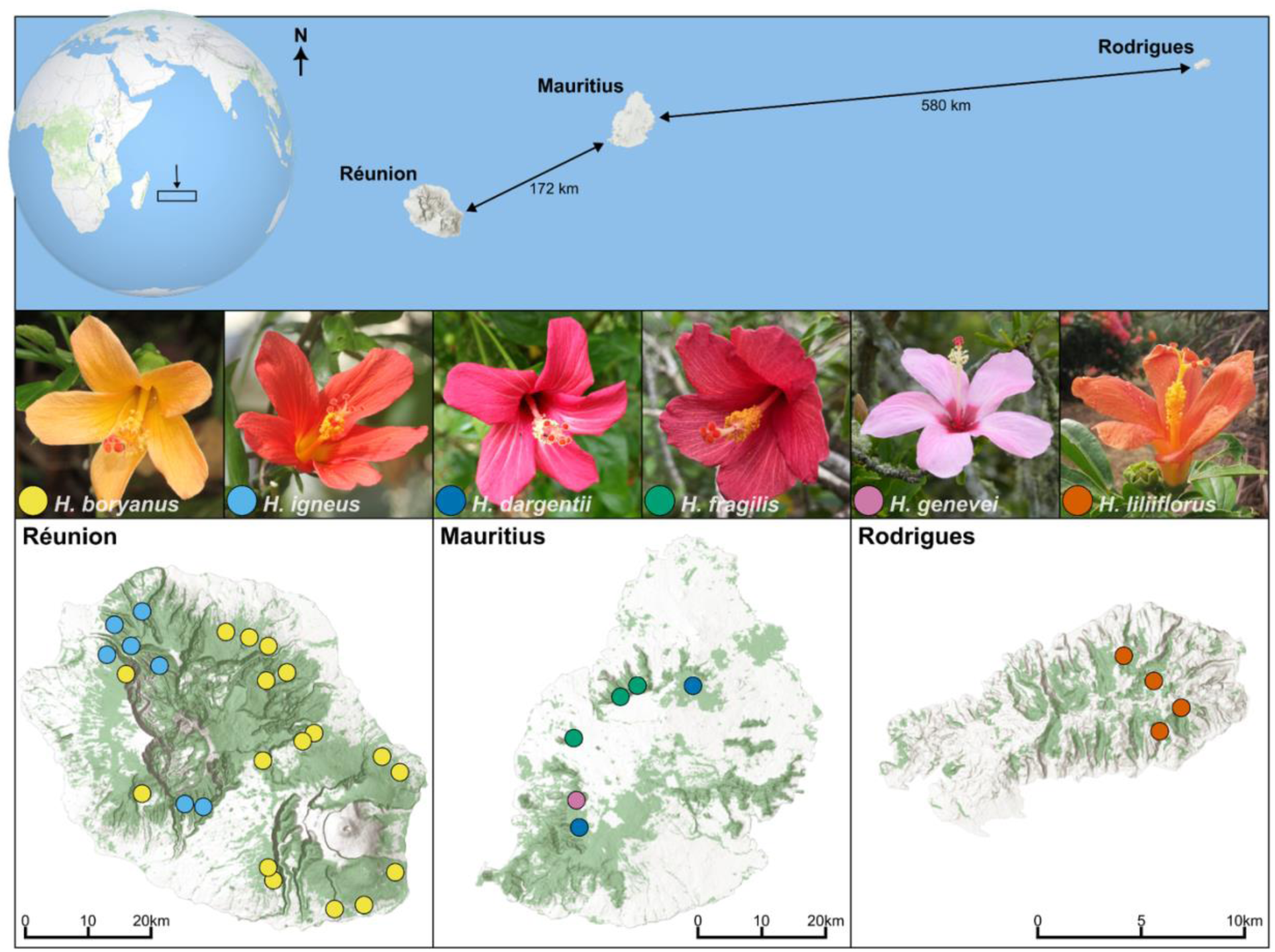
A map of the three Mascarene islands and images of the flowers of the six species occurring in the archipelago. Sampling sites in this study are indicated for each island by matching site color to the corresponding species. In Rodrigues and Mauritius, the sampled sites represent all known extant populations on each island; in Réunion, sampled sites are representative of the known distribution of each species, though intervening sites are known. See Fig. S3.1 for maps with populations labelled.

### Sampling

Samples were obtained from living plants during field trips in Mauritius and Rodrigues in March of 2018 and Mauritius and Réunion in April of 2022. Young leaves were taken from plants and immediately placed in silica gel until DNA extraction. One or more herbarium collections were made per population; duplicates are deposited at MO and MAU. We sampled 10 individuals of *H. liliiflorus* from Rodrigues and 10 individuals of *H. genevei* from the only known population in Mauritius. For *H. dargentii*, we collected five individuals from Tamarind Falls; however, only one of these Tamarind Falls individuals is known to be wild, while the other four are clones of now-dead wild individuals that once occurred there. It is not clear if these clones include all the genotypes that once occurred at Tamarind Falls, so we also sampled six additional clones of the Tamarind Falls population that are planted at an ex-situ nursery (Pigeon Wood). Our sampling of *H. dargentii*, therefore, includes 14 wild samples from Gaulettes Serrées and 11 samples of genotypes originally from Tamarind Falls. For simplicity, we refer to the populations of each species based on their relative geographic position within the island (i.e., Gaulettes Serrées = *H. dargentii* North; Tamarind Falls = *H. dargentii* South). For *H. fragilis*, we included all six individuals remaining on the nearby Letard and Ory Mountains (*H. fragilis* North) and seven collected from Corps de Garde (*H. fragilis* South), totaling in 13 samples. In Réunion, we collected 36 samples of *H. boryanus* from 13 populations and 21 samples of *H. igneus* from eight populations. To test the monophyly of the Mascarene radiation of sect. *Lilibiscus*, we included as outgroups three samples of *H. bernieri* from Madagascar, two samples each of *H. kaute* and *H. macverryi* from the Fijian archipelago, and three samples each of *H. clayi* and *H. waimeae* from the Hawaiian islands. In summary, our dataset consisted of 13 outgroup samples from four distinct geographic regions and 115 ingroup samples: 36 *H. boryanus*, 25 *H. dargentii*, 13 *H. fragilis*, 10 *H. genevei*, 21 *H. igneus*, and 10 *H. liliiflorus*.

### Library preparation and sequencing

We extracted genomic DNA from 20-30 mg of silica dried leaf samples using a modified CTAB approach (Doyle & Doyle 1987). We incorporated a preliminary sorbitol-based cleaning step following Inglis et al. (2018), using three sorbitol washes to remove mucilage. For genotyping, we employed a 2bRADseq library preparation approach (Wang et al. 2012) following previously published protocols (Mashburn et al 2023). This protocol uses a type-IIB restriction enzyme (BcgI) to cleave genomic DNA into 36 bp fragments followed by adaptor ligation, PCR amplification, sample cleanup and quantification, and pooling. Libraries were sequenced on an Illumina HiSeq 4000 using 50 bp single-end reads at Northwestern University’s NUSeq Core.

### Locus assembly and data filtering

Raw reads were inspected with FastQC (Andrews 2010) and demultiplexed using a custom script. Raw sequence reads were filtered using FASTX-Toolkit version 0.0.14 (Hannon 2010) with the settings -Q 33 -q 20 -p 90. Sequences were then assembled *de novo* using ipyrad version 0.9.84 (Eaton & Overcast 2020). Reads were clustered using a 95% similarity threshold, and clusters with <10 and >8000 reads were removed. We produced three assemblies with varying missing data thresholds based on the requirements of downstream applications. Given that maximum likelihood phylogenetic reconstruction methods are generally robust to missing data and perform better with more loci (Eaton et al. 2017), we produced an assembly that included all ingroup and outgroup samples (hereafter referred to as MascPhy_all) and retained loci found in at least 15% of samples. For species tree reconstruction in Tetrad, which is computationally intensive and thus requires a smaller sample size, we created an assembly with only ingroup samples, subsampling four samples from species in which only one population remains *(H. liliiflorus* and *H. genevei*) and eight samples from the remaining species (which each have two or more populations) and requiring loci to be found in at least 50% of all samples (hereafter referred to as MascPhy_sub). For population genomic analyses and demographic history modeling, which are more sensitive to missing data (Chattopadhyay et al. 2014; Hodel et al. 2017), we produced a strictly filtered dataset (hereafter referred to as MascStr_in), containing all ingroup samples and retaining loci found in at least 80% of samples.

### Phylogenetic reconstruction

We conducted maximum-likelihood (ML) phylogenetic analyses using IQ-TREE version 2.2.0.3 (Minh et al. 2013) with the built-in ModelFinder (Kalyaanamoorthy et al. 2017) to select the optimal nucleotide substitution model. Branch supports were assessed using the ultrafast bootstrap approximation (UFBoot2) (Hoang et al. 2018) and the Shimodaira-Hasegawa approximate likelihood ratio test (SH-aLRT) (Guindon et al. 2010; Shimodaira & Hasegawa 1999), both with 1000 replicates. Ultrafast bootstrapping (UFBoot) is more unbiased compared to normal bootstrapping, so higher UFBoot values are necessary to demonstrate a well-supported clade (Minh et al. 2013). In addition, SH-aLRT is more robust than bootstrapping methods for short branch lengths, though bootstrapping methods sample more topologies than the nearest-neighbor-interchanges method performed in the SH-aLRT test (Guindon et al. 2010). Therefore, we treated branches as well supported if they had both a UFBoot support value >= 95% and a SH-aLRT support value >= 80% (Guindon et al. 2010; Minh et al. 2013). The tree was rooted on *H. bernieri* from Madagascar, as previous phylogenetic work on the genus indicates it is sister to the rest of sect. *Lilibiscus* (Hanes et al. 2024).

We also inferred a species tree under the multispecies coalescent model using the program Tetrad, as implemented in the ipyrad-analysis toolkit (Eaton & Overcast 2020). Tetrad employs a similar algorithm as the program SVDQuartets (Chifman & Kubatko 2014) but is adapted to RAD-seq data as it resolves quartet trees using unlinked SNPs and uses bootstrap replicates to assess node support. Tetrad was run on our subsampled phylogenetic assembly (MascPhy_sub) without species assignments (all samples were tips on the tree). We explored all possible quartets (91,390) with 100 bootstrap replicates. We displayed the tree with species/populations binned into a single tip to show simplified relationships for comparison. Tetrad and ML trees were visualized using the R packages ‘ape’ (Paradis & Schliep 2019), ‘treeio’ (Wang et al. 2020), and ‘ggtree’ (Yu et al. 2017).

### Population genetic structure

We analyzed patterns of genetic structure with the MascStr_in dataset using principal component analysis (PCA), uniform manifold approximation and projection (UMAP), and the Bayesian model-based clustering program STRUCTURE version 2.3.4 (Pritchard et al. 2000). PCA and UMAP analyses used unlinked SNPs and were performed and visualized with the ipyrad-analysis toolkit (Eaton & Overcast 2020). We also examined genetic structure among all Mascarene species and separately for the species from Mauritius. The ipyrad toolkit also allows missing data to be imputed by sampling genotype frequencies within user-defined groups of samples (‘samples’ method), or by randomly sampling genotypes in k-means cluster-generated populations (‘kmeans’ method). Given the strength of species assignments shown in the ML tree, we used the ‘samples’ method in our PCA of the whole dataset to impute missing data from other samples of the same species. Then, for our PCA of the three Mauritius species, we used the ‘kmeans’ method at k = 5 to show results based on samples being grouped as five populations (2 *H. dargentii*, 2 *H. fragilis*, 1 *H. genevei*).

We also employed UMAP, a non-linear dimension reduction algorithm that seeks to find a low dimensional embedding to preserve the essential topological structure of the dataset (McInnes et al. 2018). Since user defined UMAP parameters influence how tightly samples cluster, it should not be interpreted as a projection of how closely related samples are, but instead how closely groups cluster compared to the rest of the samples in the dataset.

To assign samples to one or more genetic clusters, we ran STRUCTURE on the assembly of all ingroup samples (MascStr_in) using an admixture model with a burn-in of 250,000 generations and a run length of 500,000 generations for K values between 1 and 10 with 10 replicates per K. Results of each STRUCTURE run were analyzed in STRUCTURE HARVESTER version 0.6.94 (Earl & vonHoldt 2012), where the optimal K value was selected based on where the log likelihood values plateau (Pritchard et al. 2000), though we also examined results using the delta K method suggested in Evanno et al. (2005). Replicate runs at each K were summarized in CLUMPP (Jakobsson & Rosenberg 2007) and we removed runs that did not converge with the major cluster found by CLUMPP.

### Population genetic diversity

We calculated several population genetic metrics to further assess genetic variation and divergence among species and intraspecific populations using the assembly of all ingroup samples (MascStr_in). We first performed interspecific comparisons with samples assigned to each species. Second, we performed comparisons with samples of *H. dargentii* and *H. fragilis* separated into their disjunct populations. To produce comparable populations in Réunion, we subsetted samples into two groups, north and south, for the two Réunion species, *H. boryanus* and *H. igneus*. To make these groups, we used the 12 northernmost and 9 southernmost collections of *H. boryanus* and the six northernmost and six southernmost collections of *H. igneus*, leaving out the remaining samples. The distance between the two *H. boryanus* populations was 30.2 km and 28.7 km between the *H. igneus* populations. In Mauritius the two *H. dargentii* populations were 24 km apart and the two *H. fragilis* populations were 7.8 km apart. We used the R packages adegenet (Jombart & Ahmed 2011; Jombart & Bateman 2008) and hierfstat (Goudet 2005) to calculate Weir and Cockerham’s (1984) and Nei’s (1987) pairwise F_ST_. We also calculated expected heterozygosity (H_e_), observed heterozygosity (H_o_), and inbreeding (F_IS_) for each species and then for each subset population of interest.

### Demographic modeling

To assess the relative divergence times between species and intraspecific populations in the Mascarenes, we performed demographic modelling using Approximate Bayesian Computation (ABC) in the program DIYABC-RF (Collin et al. 2021). Given the number of species (6) and populations (10) being sampled, the number of theoretically possible scenarios was too great to test and process all of them. As individual populations in *H. boryanus* and *H. igneus* were not strongly diverged (see Results), we focused our analyses first on assessing species divergence, then population divergence in Mauritius. We tested scenarios hierarchically, with the best-fit scenario in each step informing the scenarios in the next step of modeling. In step 1 we tested six scenarios to determine the order of divergence for the six Mascarene *Hibiscus* species (Fig. S2.5). In step 2 we tested six scenarios to determine the relative timing of the divergence between populations within the Mauritian species *H. dargentii* or *H. fragilis* in comparison to the divergence between the Reunion species *H. boryanus* and *H. igneus* (Fig. S2.7). In both steps we ran 10,000 training simulations (∼1,666 simulations per model) since we were primarily interested in model choice. During the training dataset simulation, we used the default priors of a uniform distribution from 10 to 10,000 individuals for each population size (*N*) and a uniform distribution of 10 to 10,000 generations for each timeframe (*t*). We then estimated the best-fit models in steps 1 and 2 using all 10,000 training sets, five noise variables (default), 1000 out-of-bag testing samples (default), and 1000 trees in the random forest analysis. The best-fit model was the one selected by the greatest number of trees in the forest, a process which was informed by incorporating values of *d* linear discriminant analysis (LDA) axes, where *d* is equal to the number of input models minus one (Pudlo et al. 2016).

## 3.5 Results

### Sequencing, assembly, and filtering

After initial filtering of low-quality reads, the full dataset of 128 samples contained 366.06 million reads (138,592–6,567,200 per sample; average: 2,859,871). The filtered concatenated phylogenetic dataset including all 128 samples (MascPhy_all) contained 15,551 loci, 14,854 variable sites, and 50.94% missing data in the SNP matrix. The reduced phylogenetic assembly of 40 samples (MascPhy_sub) contained 10,760 loci, 7,103 variable sites, and 20.51% missing data in the SNP matrix. The strictly filtered assembly with 115 ingroup samples (MascStr_in) contained 4,878 loci, 3,529 variant sites, and 8.69% missing data in the SNP matrix.

### Phylogenetic reconstruction

ModelFinder in IQ-TREE selected HKY+F+I+R3 as the best-fit nucleotide substitution model. The ML tree showed strong UFBoot and SH-aLRT support for most interior nodes (Fig. 3.2A). Samples from Hawaii and the South Pacific formed two monophyletic groups that were placed as successive sisters to the Mascarene samples. The six Mascarene species formed a strongly supported monophyletic group.

**Figure 3.2.**
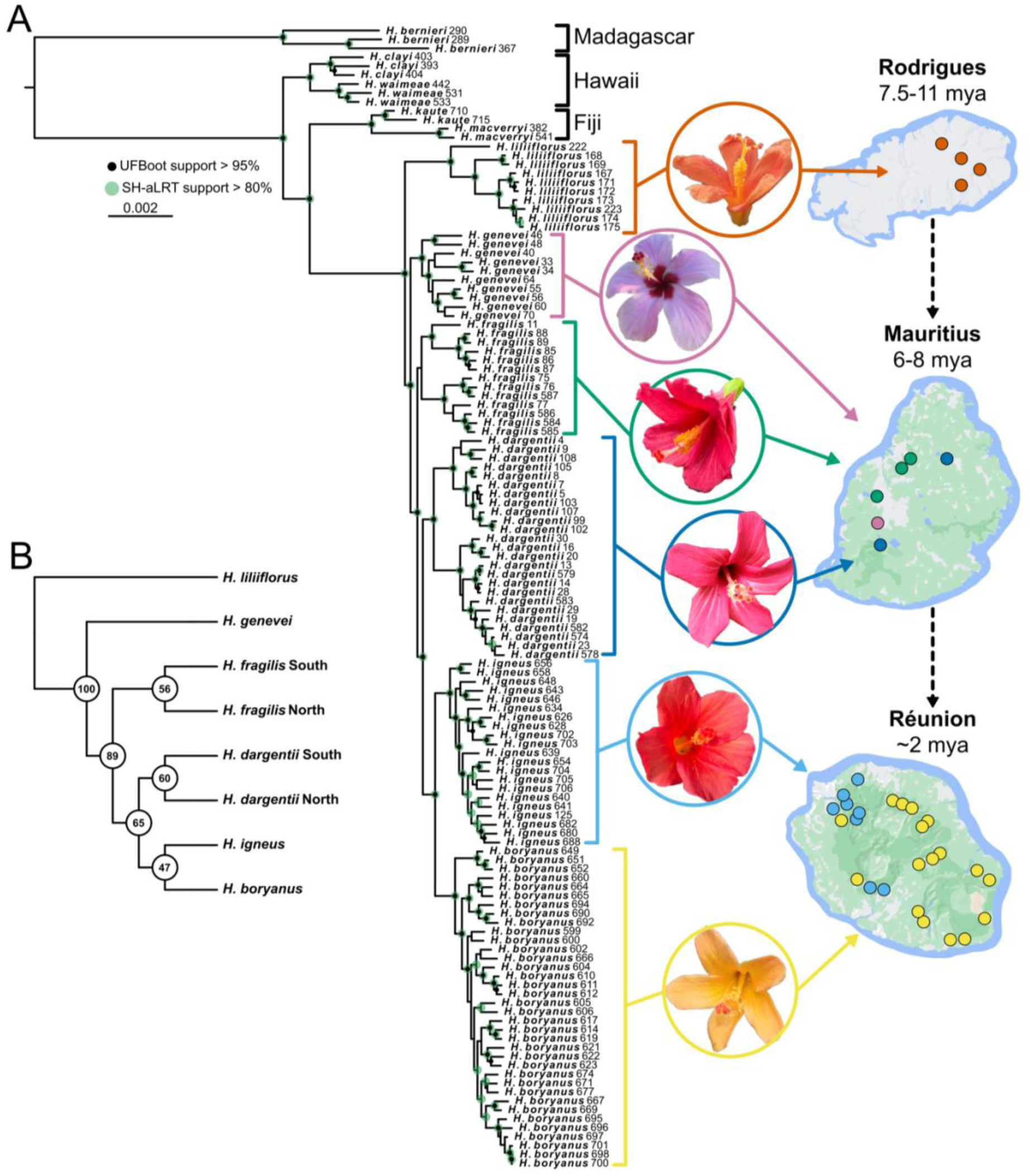
A–Maximum likelihood phylogeny of all samples including outgroups. The Mascarene Hibiscus sect. Lilibiscus form a monophyletic group with respect to outgroup species, and each of the six Mascarene species is monophyletic with strong support. Nodes with ultrafast bootstrap support > 95% after 1000 replicates are shown with a black circle. Nodes with SH-aLRT support > 80% after 1000 replicates are shown with a green circle. Scale bar indicates number of substitutions per site. Ages under island names indicates the estimated ages of the island. B–Cladogram summarizing the Tetrad majority rule consensus tree. In B species and/or populations of interest are binned to reveal simplified relationships, though the trees were produced with more samples.

Each of the six Mascarene species formed a strongly supported monophyletic group in the ML tree on relatively short branches (Fig. 3.2A). The Rodrigues-endemic *Hibiscus liliiflorus* occurred on the longest branch and was sister to the rest of the group. The three Mauritian species did not form a single monophyletic group, but instead *H. genevei* and *H. fragilis* were placed as successive sisters to *H. dargentii*. Intraspecific populations of *H. fragilis* and *H. dargentii* were resolved as distinct, strongly supported clades within each species. The internal node separating the two populations within *H. fragilis* was the only deeper node in the ML tree that did not show high support values of either UFBoot or SH-aLRT. *Hibiscus boryanus* and *H. igneus* from the youngest island of Réunion formed a monophyletic group sister to *H. dargentii* from Mauritius.

The TETRAD majority-rule consensus species tree was concordant with the species relationships in the ML tree (Fig. 3.2B, S3.2). The tree showed *H. liliiflorus* and *H. genevei* to be sister to the remaining Mascarene species with 100% bootstrap support. However, relative to the ML tree, the branch subtending *H. genevei* showed lower support values, indicating discordance between individual gene (SNP) trees in the analysis.

### Population genetic structure

STRUCTURE results generally corresponded to the species relationships found in the phylogenetic analysis with minor exceptions (Fig. 3.3A, S3.3). There was no clear consensus on the optimal K value between the Evanno or delta K method. The log probability of the data did not clearly plateau, and the greatest value of lnPK was at K = 8 (Fig. S3.4). The greatest delta K value was 2 (Fig. S3.4), a result that is often recovered as an artifact of the Evanno method (Janes et al. 2017). Figure 3.3 shows the STRUCTURE plot at K = 6, the expected value given the number of species, and K = 8, the best K according to the method of Pritchard et al. (2000). At K = 6, the only species not clearly resolved was *H. fragilis*, which was presented as potentially admixed between *H. genevei* and *H. igneus* (Fig. 3.3A). However, this history was not supported at K = 8, nor does it seem biologically feasible as the species occur on different islands. At K = 8, some limited gene flow and introgression is visible between to the two Réunion species, *H. igneus* and *H. boryanus*, though no recent F1 or F2 hybrids were recovered.

**Figure 3.3.**
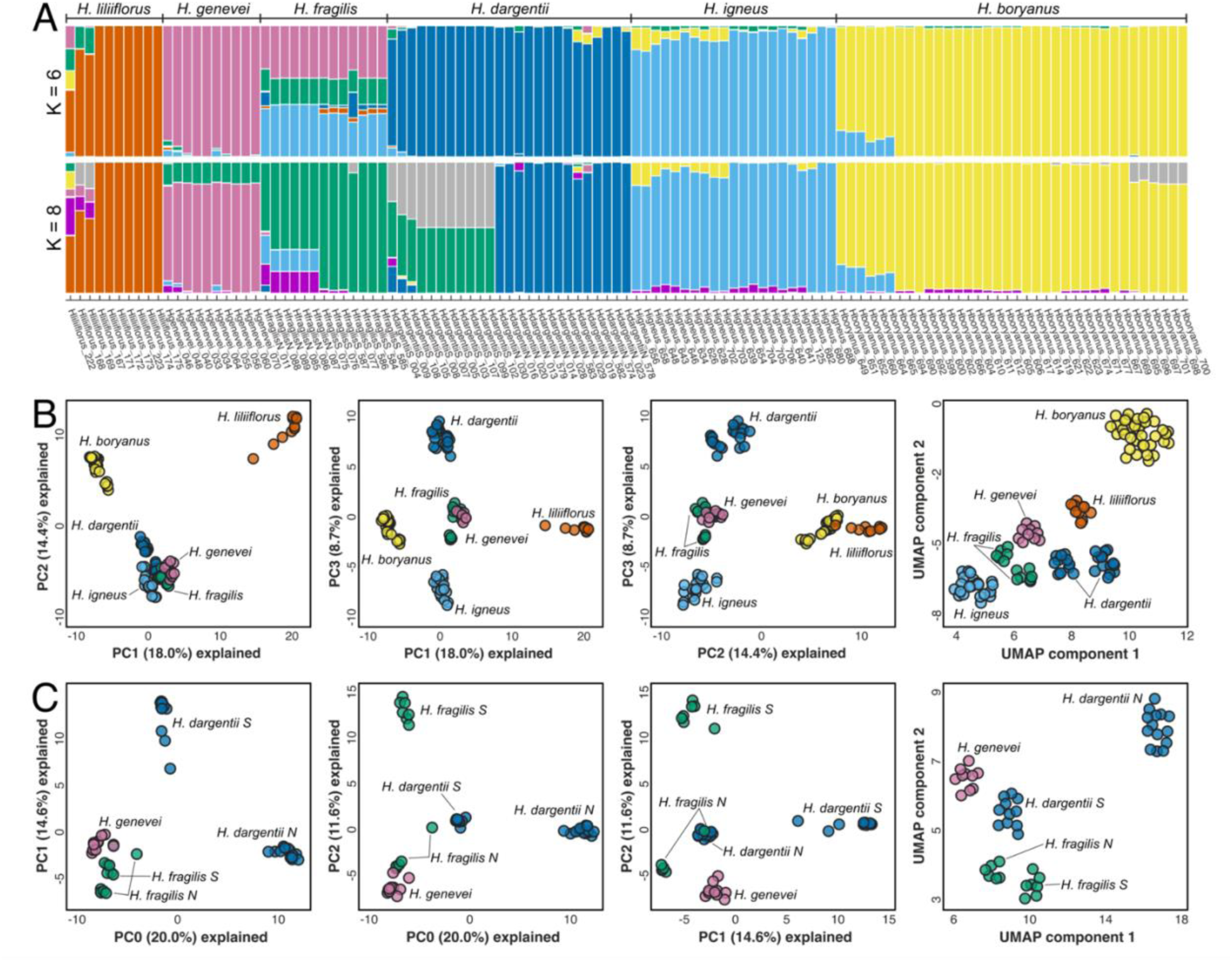
A–STRUCTURE plots at K=6, showing relationships assuming six genetic clusters, and K=8, assuming 8 genetic clusters. Samples from the northern and southern population of each species can be identified by the N and S in the sample name at the bottom of the plot. Colors represent the genetic groups, lines represent individuals, with the individuals assigned to the same color indicating individuals that are related. B–PCA plots showing the first three PCs and UMAP plot with samples of all six species. C–PCA plots showing the first three PCs and UMAP plot with samples of the three species in Mauritius; northern and southern populations of *H. dargentii* and *H. fragilis* are indicated.

When examining relationships at K = 8, STRUCTURE differentiated the two populations within each of *H. fragilis* and *H. dargentii* (Fig. 3.3A). At this K value, neither population of *H. fragilis* is shown as having equal proportions of *H. genevei* and *H. igneus* ancestry, and the southern population consists entirely of one ancestry group. *Hibiscus genevei* is shown to share some ancestry with *H. fragilis* (green color). For *H. dargentii*, the northern population consists of a single ancestry group, with the southern population approximately half of ancestry similar to *H. fragilis* and half an independent ancestry group (grey color).

PCA and UMAP were clearer in their support for species relationships compared to STRUCTURE. The PCA’s containing all six species showed five clusters, with *H. fragilis* and *H. genevei* occurring together (Fig. 3.3B). However, these two species were differentiated in the UMAP at a similar level to the other species. *Hibiscus liliiflorus* was the most diverged species in the PCA plots; however, in the UMAP *H. boryanus* formed the most distant cluster of samples. Meanwhile, *H. igneus* from Reunion clustered more closely with the Mauritian species than to the other species from Reunion, *H. boryanus,* in the PCA and UMAP.

In the PCA and UMAP of the three Mauritian species (Fig. 3.3C), the intraspecific populations of *H. fragilis* and *H. dargentii* showed high levels of differentiation. In the PCA plots, the northern *H. fragilis* population was consistently placed close to *H. genevei*, which corresponds to the STRUCTURE plot at K = 8 that showed the two groups have some shared ancestry. In the UMAP, the two *H. fragilis* populations we placed close to each other, and the most differentiated group was the northern *H. dargentii* population. This is also consistent with the STRUCTURE plots at K = 8, which found the northern *H. dargentii* population shared almost no ancestry with any of the other populations in Mauritius, including the southern *H. dargentii* population.

### Population genetic diversity

Across the six *Hibiscus* species, expected heterozygosity ranged from 0.0374 to 0.1015 and observed heterozygosity ranged from 0.0263 to 0.0891 (Table 3.2). Both values were highest in *H. liliiflorus,* which occurs on the oldest island. *Hibiscus genevei* from Mauritius also had relatively high rates of expected and observed heterozygosity, which likely resulted from decades of human-mediated planned outcrossing among individuals in the only extant population. Rates of heterozygosity for the range-restricted species *H. dargentii* and *H. fragilis* in Mauritius were intermediate, whereas *H. boryanus*, the species with the largest extant population size and the most widespread range in the youngest island, Réunion, had the lowest value of both measures. Measures of heterozygosity did not differ substantially for the subset populations of the four species with multiple populations compared with the species as a whole (Table 3.2).

**Table 3.2.**
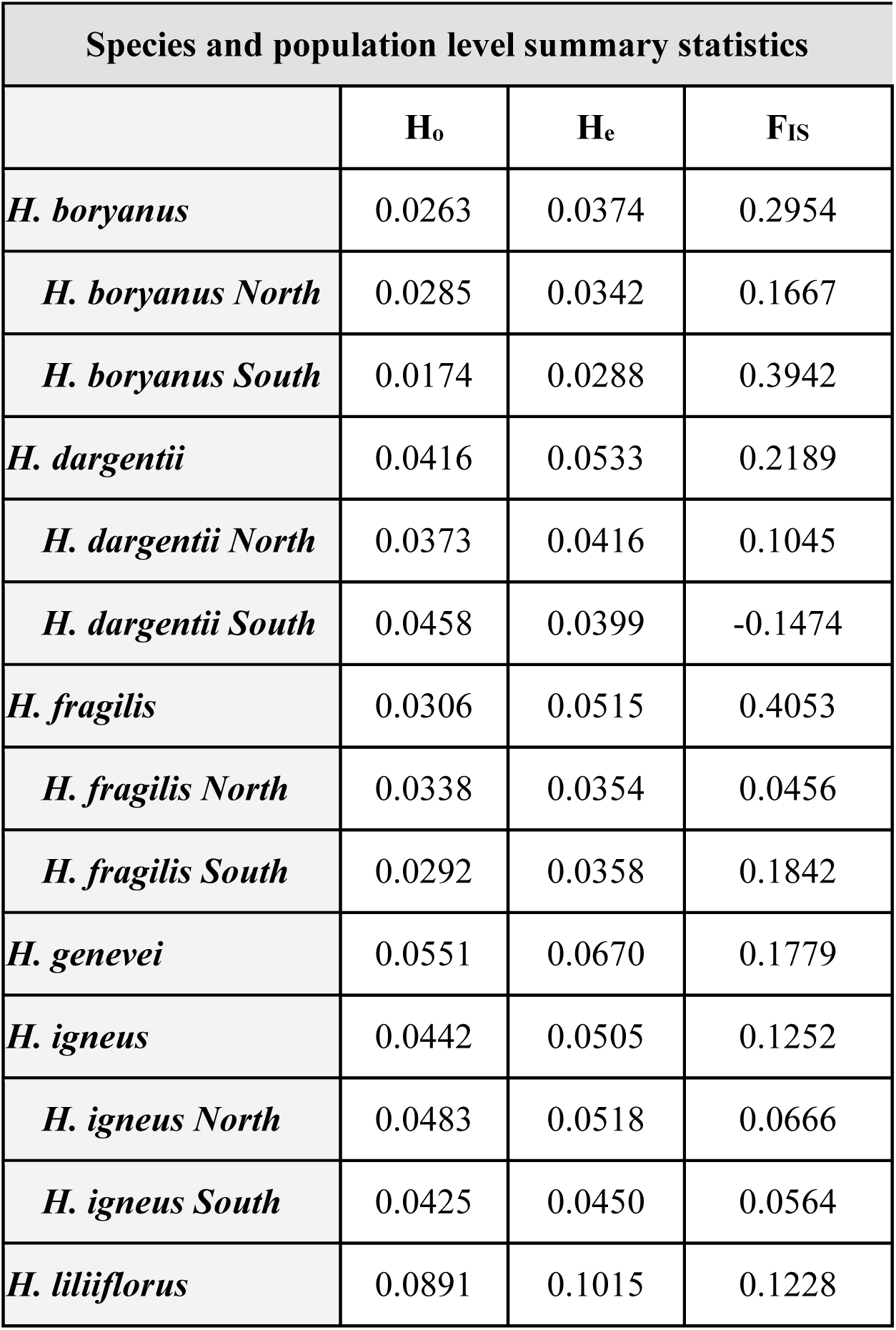
Summary population genetic statistics for each species and subset populations. The species level statistics were calculated with all samples for that species, and the population level statistic were calculated with the subset of samples from each population. For *H. dargentii* and *H. fragilis*, the two populations together comprise all of the samples found in the species; for *H. boryanus* and *H. igneus*, the two populations are made up of samples taken from the northernmost and southernmost edges of the species’ ranges.

Species-level F_IS_ values ranged from 0.1228 to 0.4053 (Table 3.2). The highest value was found in *H. fragilis*, signifying very high levels of inbreeding within the species. However, when looking separately at the two populations of *H. fragilis*, F_IS_ values fell dramatically, suggesting a Wahlund effect. The higher F_IS_ value for *H. dargentii* as a whole compared to the values for each population may also be caused by a Walhund effect. *H. boryanus* also demonstrated a very high value of F_IS_ (0.295), which is surprising because of the comparatively large population sizes of the species; inbreeding was even more severe for the southern population (0.394) which is more reduced and fragmented than the eastern and northern parts of the species’ range. In addition, populations of the two restricted Mauritian species did not demonstrate substantially higher rates of inbreeding than the two more common Réunion species.

Pairwise F_ST_ (Nei’s) comparisons between species were high (14 of 15 comparisons > 0.25) and ranged from 0.230 to 0.516 (average: 0.359; Table 3.3). *Hibiscus liliiflorus* exhibited the highest average pairwise F_ST_ value (0.456) with other species, likely due to greater time since divergence and less connectivity to the other species. The two Réunion species, though recently diverged and seemingly experiencing some limited gene flow (Fig. 3.3A), also showed high pairwise F_ST_ (0.391). The three Mauritian species had the three lowest pairwise F_ST_ values, suggesting historic connectivity among species within Mauritius.

**Table 3.3.** Pairwise F_ST_ calculations between the six Mascarene *Hibiscus* sect. *Lilibiscus* species. The bottom left shows Weir and Cockerham’s F_ST_ with average values for each species at the bottom. The top right shows Nei’s F_ST_ with average values for each species at the right.

|  | *H. boryanus* | *H. dargentii* | *H. fragilis* | *H. genevei* | *H. igneus* | *H. liliiflorus* | Nei Average |
| --- | --- | --- | --- | --- | --- | --- | --- |
| *H. boryanus* |  | 0.347 | 0.401 | 0.383 | 0.391 | 0.516 | 0.408 |
| *H. dargentii* | 0.358 |  | 0.257 | 0.251 | 0.304 | 0.442 | 0.320 |
| *H. fragilis* | 0.422 | 0.258 |  | 0.230 | 0.267 | 0.446 | 0.320 |
| *H. genevei* | 0.429 | 0.261 | 0.233 |  | 0.274 | 0.399 | 0.307 |
| *H. igneus* | 0.402 | 0.305 | 0.269 | 0.285 |  | 0.478 | 0.343 |
| *H. liliiflorus* | 0.546 | 0.432 | 0.422 | 0.360 | 0.468 |  | 0.456 |
| WC Average | 0.431 | 0.323 | 0.321 | 0.314 | 0.346 | 0.445 |  |

Pairwise F_ST_ comparisons at the population level for the four focal species revealed interesting patterns of differentiation based on island of origin (Table 3.4). Values of Nei’s F_ST_ ranged from 0.077 to 0.578 when incorporating *H. genevei* and *H. liliiflorus* into the comparisons (average: 0.425) (Table S3.1). Comparisons between the northernmost and southernmost populations of the Réunion species *H. igneus* (0.077) and *H. boryanus* (0.162) showed the lowest values (Table 3.4). In contrast, pairwise F_ST_ values between the two populations within either *H. dargentii* or *H. fragilis* from Mauritius were much higher: 0.387 and 0.472, respectively (Table 3.4) and were greater than the average Nei’s F_ST_ value from the inter-species comparisons (0.359, Table 3.3), suggesting that genetic differentiation between each population of *H. dargentii* and *H. fragilis* is on par with species-level differentiation among the clade as a whole. Notably, the F_ST_ value between the northern and southern populations of *H. fragilis*, separated by only 7.8 km, was greater than nearly all species-level comparisons except comparisons between the distantly related *H. liliiflorus* and the two Réunion species (Table 3.3).

**Table 3.4.**
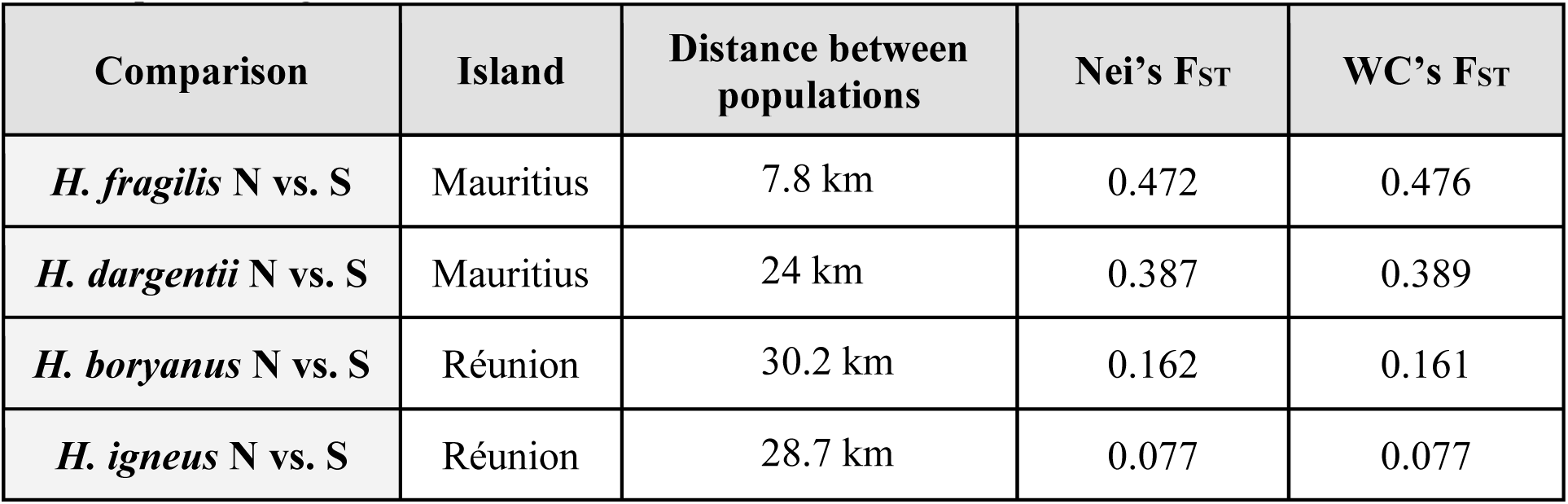
Pairwise F_ST_ calculations between north and south populations of the four focal *Hibiscus* species that have two or more populations. The northern and southern populations of *H. dargentii* and *H. fragilis* are the only populations of each species and together comprise all sampled individual. The northern and southern populations of *H. boryanus* and *H. igneus* are selected samples from the two ends of each species’ range.

### Demographic modeling

In step 1 of demographic modeling, we tested six scenarios of species-level divergence which incorporated different models of dispersal and divergence for the six species on the three Mascarene Islands (Fig. S3.5). DIYABC-RF found the best-fit model to be scenario 5 (Fig. S3.6) with 663 of 1000 trees in the random forest and a posterior probability of 0.93 (Fig. 3.4A). Scenario 5 demonstrated a stepwise process of divergence for each species concordant with the ML topology. The second-best model selected was scenario 3 with 197 trees. All other scenarios were supported by fewer than 50 trees each (Fig. S3.5).

**Figure 3.4.**
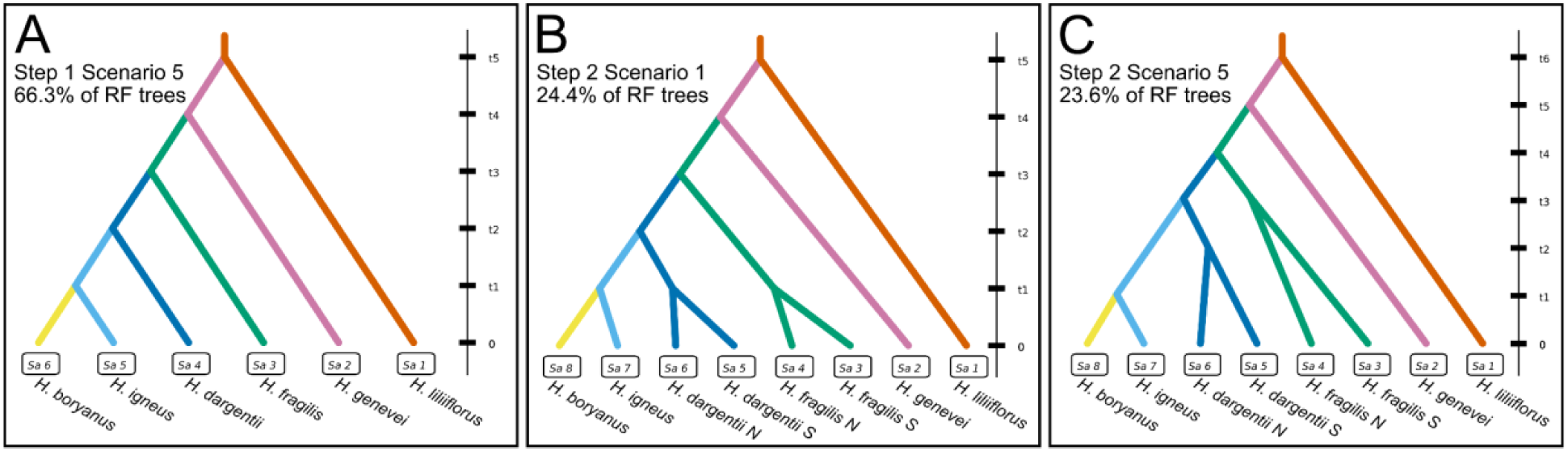
Best-fit scenarios from coalescent-based modeling in DIYABC-RF. A–In step 1, which grouped samples into six species, scenario 5 was strongly supported as the best-fit scenario, showing stepwise speciation that corresponds to the topology in the phylogenetic analyses. B & C–In step 2, which separated samples from the northern and southern populations of *H. dargentii* and *H. fragilis* into distinct groups, two scenarios received similar levels of support. In B, scenario 1 shows the populations of *H. dargentii* and *H. fragilis* diverging at the same time point (t1) as the *H. boryanus* and *H. igneous*. In C, scenario 5, the populations of *H. fragilis* split (t3), followed by the populations of *H. dargentii* (t2), followed by the split between *H. boryanus* and *H. igneus* (t1).

We used the topology from scenario 5 in step 1 to design our models in step 2, which sought to identify relative order of population and species-level divergence events. We included eight tips on each tree—one tip for each population of *H. fragilis* and *H. boryanus*, and one tip for the four other species (Fig. S3.7). Each scenario included the same stepwise speciation process as found in step 1, but varied in the relative order of 1) the divergence between the two populations of *H. fragilis*, 2) the divergence between the two populations of *H. dargentii*, and 3) the divergence between *H. igneus* and *H. boryanus* in Réunion. Because the modeled scenarios in step 2 were only slightly different from each other, DIYABC-RF found nearly equal support for two models (Fig. S3.7). The best-fit model in step 2 was scenario 1 with 244 of 1000 trees in the random forest and a 0.536 posterior probability (Fig. 3.4B). The second-best model was scenario 5, which was supported by 236 trees in the random forest (Fig. 3.4C). Scenarios 2 and 6 were supported by 186 and 164 trees, respectively, and scenarios 3 and 4 were supported by fewer than 100 trees each. Whereas scenario 1 presented a model with all three divergence events occurring simultaneously, scenario 5 modelled the divergence of the *H. fragilis* populations occurring before the divergence of the *H. dargentii* populations, followed by the two Réunion species. In fact, after scenario 1, the next three best-supported scenarios (5, 2, 6) all modelled the divergence of the two populations of *H. fragilis* as more ancient than the divergence of *H. boryanus* and *H. igneus* in Réunion (Fig. S3.7). This means that 83% of trees in the random forest supported a model where the two populations of *H. fragilis* diverged for at least as long or longer than the two species in Réunion. In addition, the top three scenarios (1, 5, 2) also modelled the divergence of the two populations of *H. dargentii* as equal to or more ancient that the divergence of *H. boryanus* and *H. igneus*.

## 3.6 Discussion

The success of the general dynamic model of oceanic island biogeography (Whittaker et al. 2008) stems from repeated studies demonstrating patterns matching its predictions. However, to our knowledge, no studies have tested the implicit prediction of the GDM that main driver of speciation shifts from ecological speciation to drift over the course of an island’s ontogeny. In this study, we comprehensively sampled populations of the six Mascarene species in *Hibiscus* sect. *Lilibiscus* to assess species boundaries, evolutionary relationships, and demographic history, which we used to assess whether patterns are consistent with the hypothesis that the evolutionary processes affecting speciation will change over the ontogeny of islands. However, for a clade occupying an island archipelago to be suitable for testing this hypothesis, it must be a monophyletic group that resulted from a single dispersal event. The relationships recovered in our ML phylogeny indicated that the Mascarene radiation is monophyletic and may have resulted from a long-distance dispersal event from the Fijian archipelago, despite the fact that the Fijian islands are approximately 16.5 times farther away from the Mascarenes than the nearest possible source of dispersers in Madagascar. However, more sampling is necessary from Madagascar to corroborate this result.

### Phylogenetic relationships resolve taxonomic questions

Taxonomic treatments of the Mascarene *Hibiscus* sect. *Lilibiscus* prior to Mashburn et al. (2024) recognized four species on the three islands: *H. boryanus*, *H. fragilis*, *H. genevei*, and *H. liliiflorus* (see Friedmann 1987). Though six distinct floral morphologies (see Fig. 3.1) were known by Friedmann (1987), three of these were treated under *H. boryanus* because all three morphologies were similar in quantitative measures of floral parts, and because of confusion over species names, which have been misapplied in the Mascarenes going back to the 19th century. Mashburn et al. (2024) addressed these issues, delimiting six morphologically distinct species, two of which—*H. dargentii* and *H. igneus*—were newly described from specimens previously treated under *H. boryanus* based on morphological and life history traits.

The phylogenomic analyses performed in the current study (Fig. 3.2) support the delimitation of six species in the Mascarenes. The Mascarene radiation likely occurred very rapidly, as evidenced by the short branch lengths in the ML tree. Such rapid radiations can lead to discordance between gene trees caused by incomplete lineage sorting, which could explain the low support values in the internal nodes of the Tetrad species tree (Fig. 3.2B). Ancestral gene flow, particularly between *H. fragilis* and *H. genevei*, may also explain the observed pattern of low internal node support in the Tetrad tree. *Hibiscus genevei* and *H. fragilis* consistently shared ancestry in STRUCTURE analysis and were often found clustering together in PCA and UMAP plots (Fig. 3.3), despite the fact that the taxa are morphologically divergent. In addition, the lowest inter-species F_ST_ values were found between those two species (Table 3.3). Such admixture, if it occurred, is likely ancient, as the pattern of shared genetic history was consistent among all individuals in each population and there was no evidence of recent gene flow between individual samples.

### Patterns of species diversity and island biogeography fit the predictions of the GDM

The relationships shown in the ML and Tetrad phylogenies (Fig. 3.2), as well as in the best-fit ABC models (Fig. 3.4), revealed a pattern of dispersal and speciation that fits the species age–island age correlation predicted by the progression rule and the GDM. The earliest diverging species in the Mascarene radiation, *H. liliiflorus*, occurs on the oldest island (Rodrigues) and is most distantly related to the rest of the radiation in the Mascarenes, shown by the long branch separating *H. liliiflorus* from the rest of the radiation in the ML tree and the highest average pairwise genetic differentiation between it and the other five species (Table 3.3). The common ancestor of *H. liliiflorus* and the remaining Mascarene taxa likely dispersed from Rodrigues to the next oldest island, Mauritius, and experienced a small radiation there. All three Mauritian species occur in small, restricted populations on the high central plateau and mountaintops. The two species in Réunion, then, share a common ancestor with *H. dargentii*, suggesting that the common ancestor of *H. dargentii* and the Reunion species dispersed from Mauritius to Réunion, the youngest island of the archipelago, where it diverged into two species.

The Mascarene radiation of *Hibiscus* sect. *Lilibiscus* also displays a hump-shaped pattern of species diversity across islands of successive ages, consistent with the GDM. Though sect. *Lilibiscus* is not particularly species-rich in the Mascarenes compared to other radiations (e.g., *Diospyros*, *Dombeya*), the observed patterns of species richness fit the predictions of the GDM, with one species in the oldest island, three on the middle-aged island, and two recently diverged species on the youngest island. Notably, the observed pattern was not driven by multiple events of inter-island dispersal and vicariance, but instead by in-situ speciation on individual islands.

The fact that patterns of both phylogenetic relationships and species richness patterns fit with the predictions of the GDM would suggest that the Mascarene *Hibiscus* sect. *Lilibiscus* radiation evolved under a scenario where the group arrived in Rodrigues before the emergence of Mauritius and Réunion. This conclusion fits with recent dating of sect. *Lilibiscus* that places the stem age of the group at ∼14.66 mya, older than the estimated ages of the uplift of Rodrigues (∼7.5-11 mya).

### Island ontogeny predicts drivers of diversification

The Mascarene radiation of *Hibiscus* sect. *Lilibiscus* exhibits rapid shifts in floral morphology, especially in the diversity of color combinations (Fig. 3.1), suggesting that speciation may have been driven by the group’s ability to rapidly adapt to a diversity of pollinators, most likely birds. However, unfortunately, the majority of these bird species have long been extinct (Cheke 2013; Cheke & Hume 2008), making studies of pollinator associations difficult. However, our population-level sampling of all six species provides some insights into the drivers of adaptation and speciation in the group and suggests a relationship between speciation processes and island ontogenetic stage.

Genetic differentiation between *H. boryanus* and *H. igneus* in Réunion did not entirely fit our expectations, given that the two species are recently diverged sister species, which may be the result of ecological speciation. PCA and UMAP clustering showed *H. boryanus* to be strongly diverged from the rest of the group (Fig. 3.3B) whereas *H. igneus* clustered more closely with the species from Mauritius. In addition, *H. boryanus* showed greater average pairwise F_ST_ differentiation than the remaining four species and diverged from *H. igneus* at a similar level to the species in Mauritius (Table 3.3). *Hibiscus boryanus* and *H. igneus* have partitioned the island of Réunion along the strong windward/leeward rainfall and temperature gradient that occurs in Réunion, a pattern that has been recovered in other systems studied on the island (Garot et al. 2018). *Hibiscus boryanus* occurs on the wetter, windward side of Réunion, a habitat which has selected for a greater proportion of vertebrate-dispersed fleshy-fruiting species compared to other environments (Albert et al. 2018, 2021). We hypothesize that *H. boryanus* has experienced strong ecological selection in Réunion, leading to greater genetic differentiation from closely related species compared with its sister species, *H. igneus*. This is evidenced by some unique morphological and life history traits seen in *H. boryanus*, particularly a distinct flowering season with a semi-deciduous habit where leaves drop at the onset of flowering, and fleshy fruits that drop to the ground with seeds maturing inside the fruit (Mashburn et al. 2024). All other Mascarene species flower throughout the year and retain their leaves, and all other species in the global distribution of sect. *Lilibiscus* (∼30 species) have fruits that are dry capsules which dehisce on the plant allowing individual seeds to drop to the ground. The relatively rapid evolution of such novel traits in *H. boryanus* could account for the observed strong genetic differentiation between *H. boryanus* and the rest of the Mascarene species. These patterns may also be consistent with the hypothesis that the ecological speciation may be an important evolutionary force on younger islands.

In contrast to the potential ecological speciation that occurred in Reunion, population-level differentiation within *H. dargentii* and *H. fragilis* are more suggestive of genetic drift acting on species occurring on the island of Mauritius. Values of genetic differentiation between populations in the Mauritian species were similar or higher than those found in comparisons among species (Table 3.3), whereas much lower population divergence was found between more geographically distant populations in *H. boryanus* and *H. igneus* in Reunion. In addition, values of observed heterozygosity and inbreeding within the populations in Mauritius were not elevated compared to the other species, indicating that this pattern was not driven by recent population-level process inducing inbreeding (Table 3.2). Instead, coalescent modeling in DIYABC-RF found that the divergences between the populations of *H. dargentii* and *H. fragilis* are likely as old or older than the speciation event separating *H. boryanus* and *H. igneus* (Fig. 3.4). Yet, the populations within *H. dargentii* and *H. fragilis* are morphologically indistinguishable and occupy similar habitats, so neither morphological nor environmental local adaptation is apparent.

We suggest that the observed strong genetic divergence between intraspecific populations of *H. dargentii* and *H. fragilis* is a result of vicariance and drift. We hypothesize two non-exclusive scenarios that could drive this pattern. First, all three Mauritian species occur in higher elevations, which may have been more widespread when the island was younger. As the island subsided, the extant populations could be the last remnants of communities remaining small patches of suitable high-elevation habitat. Alternatively, all three species occur on formations consisting of the oldest (earliest) exposed series of volcanic activity. The species could have been more widespread at an earlier time, but other populations were wiped out by younger (later) series of lava flows. This second hypothesis, though, does not explain why the species were not able to recolonize habitats after younger series of volcanic activity. In either case, it is evident that, though the radiation of the Mascarene sect. *Lilibiscus* in Mauritius and Réunion likely proceeded rapidly, the divergence of populations of *H. dargentii* and *H. fragilis* in Mauritius occurred soon after the advent of the species, and the presence of relict populations is not apparently the product of recent human-mediated processes such as habitat loss occurring on the island.

### Conservation implications and future directions

In light of the extensive deforestation that has taken place in the Mascarenes over the last 300 years, especially in Rodrigues and Mauritius, the conservation community has largely assumed that the rarity of *H. dargentii*, *H. fragilis*, *H. genevei* and *H. liliiflorus* has been driven by habitat loss, and that the few remaining populations are relicts of a more widespread range for each species. In Rodrigues, for example, botanical collections began only after the island was almost entirely deforested (Balfour 1879), and thus collections of *H. liliiflorus* on the island largely come from relict individuals across the island.

In contrast to Rodrigues, botanical explorations in Mauritius and Réunion took place concurrent with deforestation. In Mauritius, *H. dargentii* and *H. genevei* have only ever been collected or reported from the extant populations. Only *H. fragilis* has been reported from ∼4 additional mountaintop populations along the western edge of the island; these populations have not been collected in recent times and may now be extirpated (Mashburn et al. 2024). The implication of these observations is that each species may be naturally rare, which may be exacerbated by human-mediated habitat loss.

In this study, we aimed to assess the evolutionary relationships and drivers of diversification in the Mascarene *Hibiscus* sec. *Lilibiscus*. This study is the first, to our knowledge, to demonstrate the presence of shifting drivers of divergence over the course of an island’s life span, fitting with the complex predictions of the GDM. The patterns of genetic differentiation found in this study are consistent with the hypothesis that ecological speciation has driven the most recent speciation event on the youngest island of Réunion, whereas divergence via genetic drift appears to be a predominant force affecting species on the middle-aged island of Mauritius. Although not tested here, it is likely that extinction is an important force affecting species the oldest island, Rodrigues, which is in a late stage of subsidence. However, future research on the group is necessary to conclusively demonstrate that local adaptation to environmental gradients is driving speciation in Réunion. In the same vein, similar population-level analyses are necessary for groups that span the three Mascarenes islands, as well as other archipelagos, to investigate whether these many taxa show evidence of similar forces driving radiations in insular systems across the globe.

## 3.7 Acknowledgements

This project would not have been possible without the generous support of a large team of people. We would like to thank all those who joined us in the field, shared insights and information, and helped acquire collection and export permits. In Mauritius and Rodrigues: Vikash Tatayah, Reshad Jhangeer-Khan, Alfred Bégué, Kerseley Pynee, Stiven Desbleds, Richard Peerun. In Réunion: Timothée Le Péchon, Lucile Jourdain Fievet, Jean Maurice Tamon, Joel Dupont, Dominique Oudin, Arnaud Rhumeur, Alexis Gorissen, Bertrand Mallet. All field work and sample transport was performed under the following permits: Mauritius and Rodrigues – NPCS NP 46/3 V1 and MBG-CBD-MAD PIC 2015-1; Réunion – CBN-CPIE Mascarin 34067135300035.

## 3.9 Data Availability

Morphometric data and scripts for data analysis and production of figures will be available on GitHub (github.com/brockmashburn) upon publication of the manuscript in a peer reviewed journal unless the journal of publication requires these materials to be made available elsewhere.

## 3.10 Supplementary Materials

**Figure S3.1.**
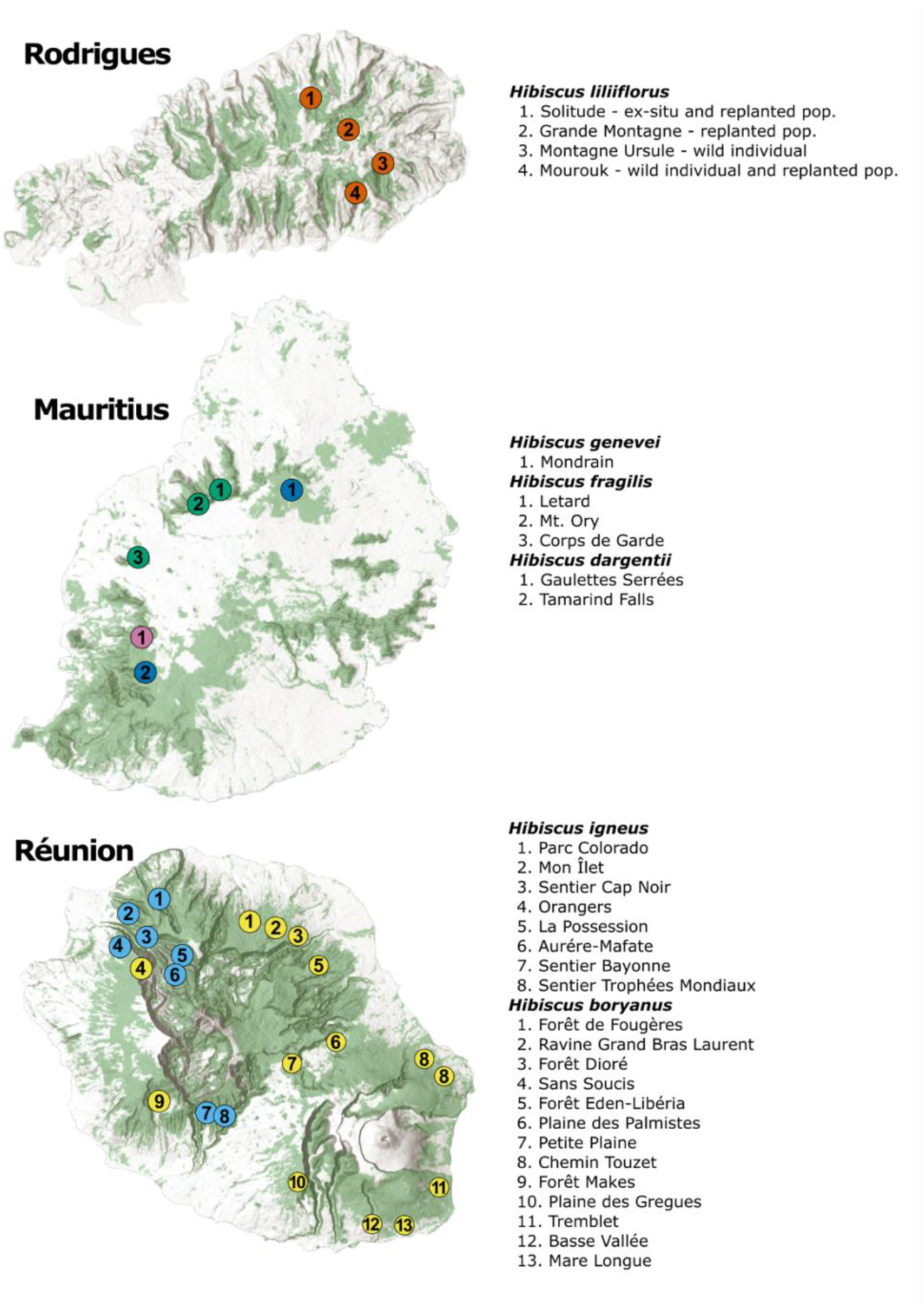
Maps of each island showing sampling locations included in this study with the names of each location to the right. The names of each sampling site correspond to Table 3.1. Green shading on each island indicates existing forest cover but does not indicate areas of native/primary forest.

**Figure S3.2.**
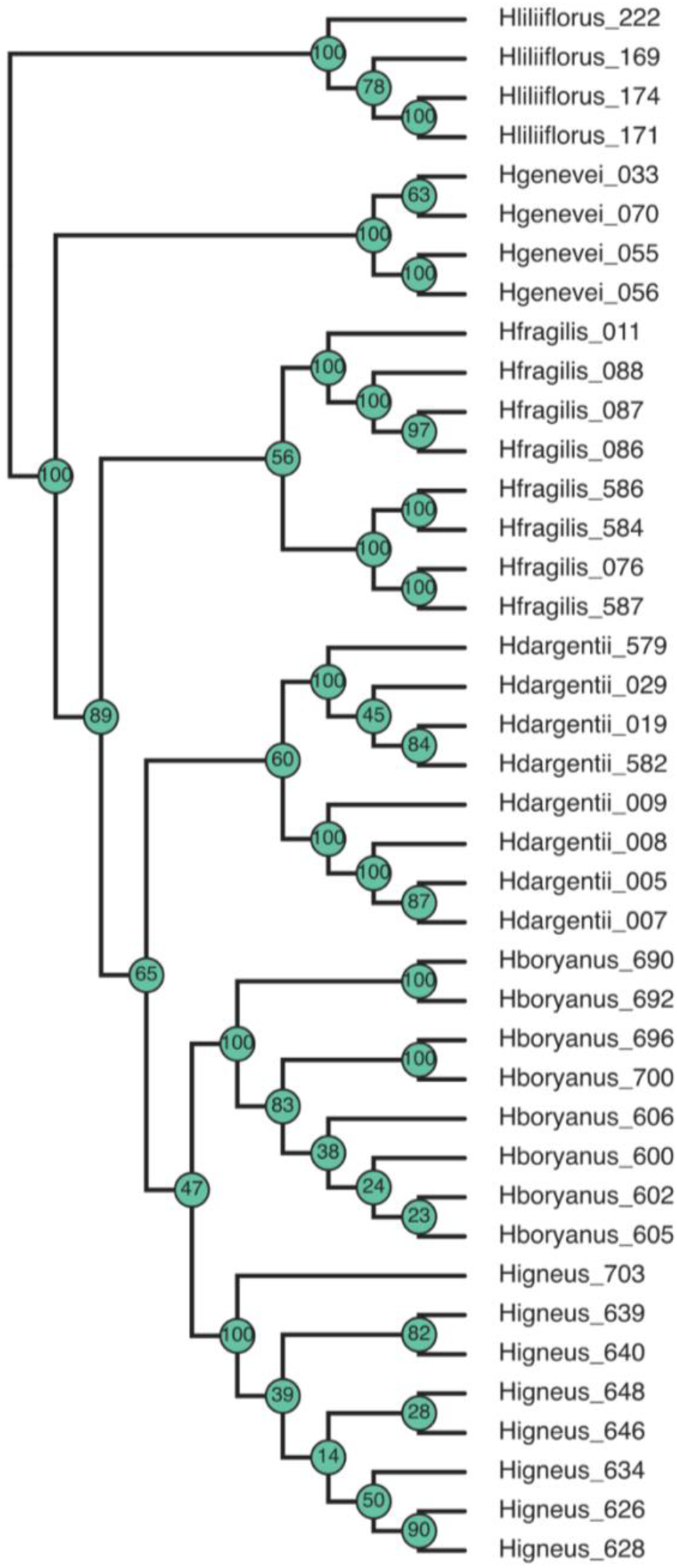
The Tetrad majority rule consensus tree showing all the tips and relationships between individual samples.

**Figure S3.3.**
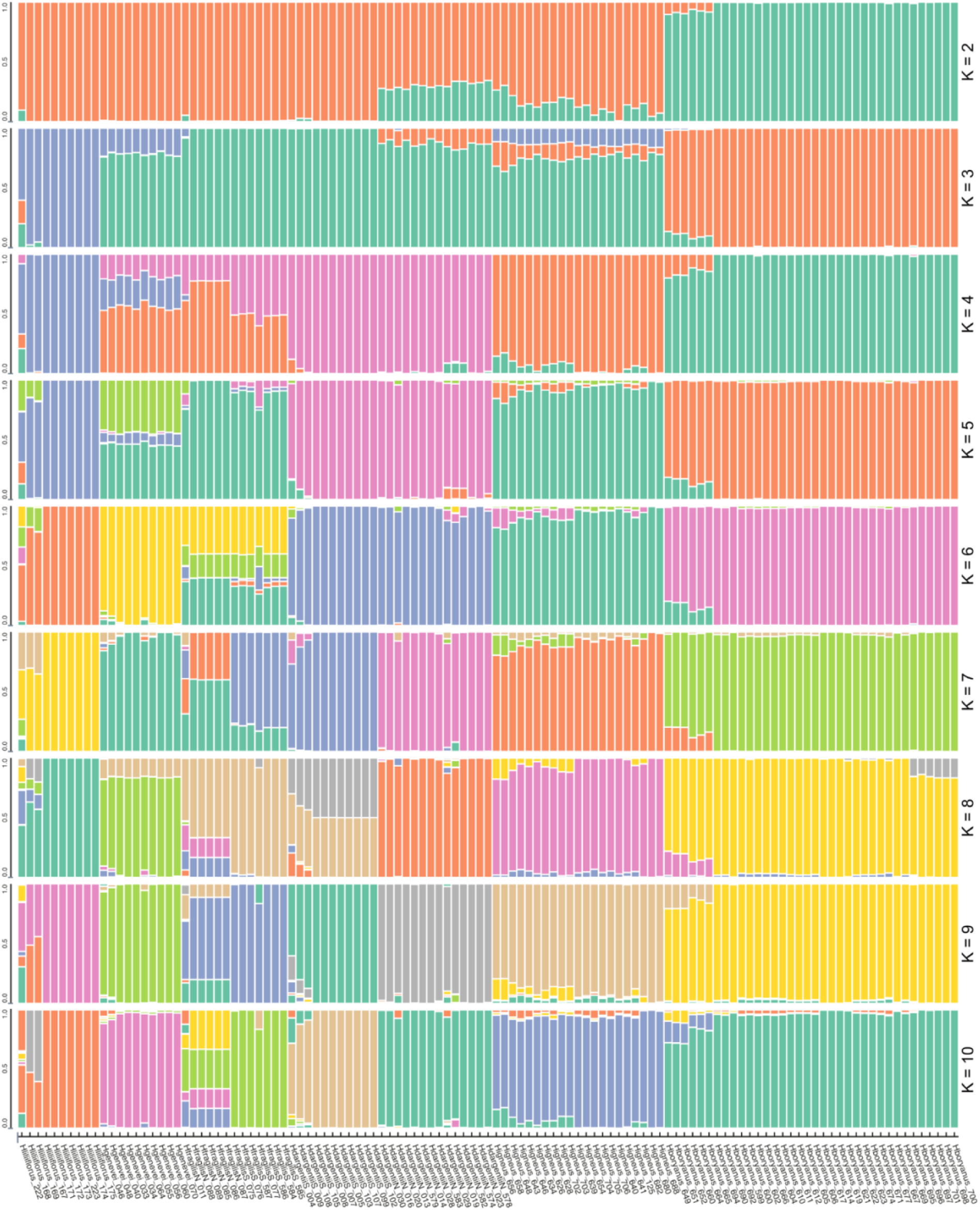
STRUCTURE plots of all tested K values (2-10).

**Figure S3.4.**
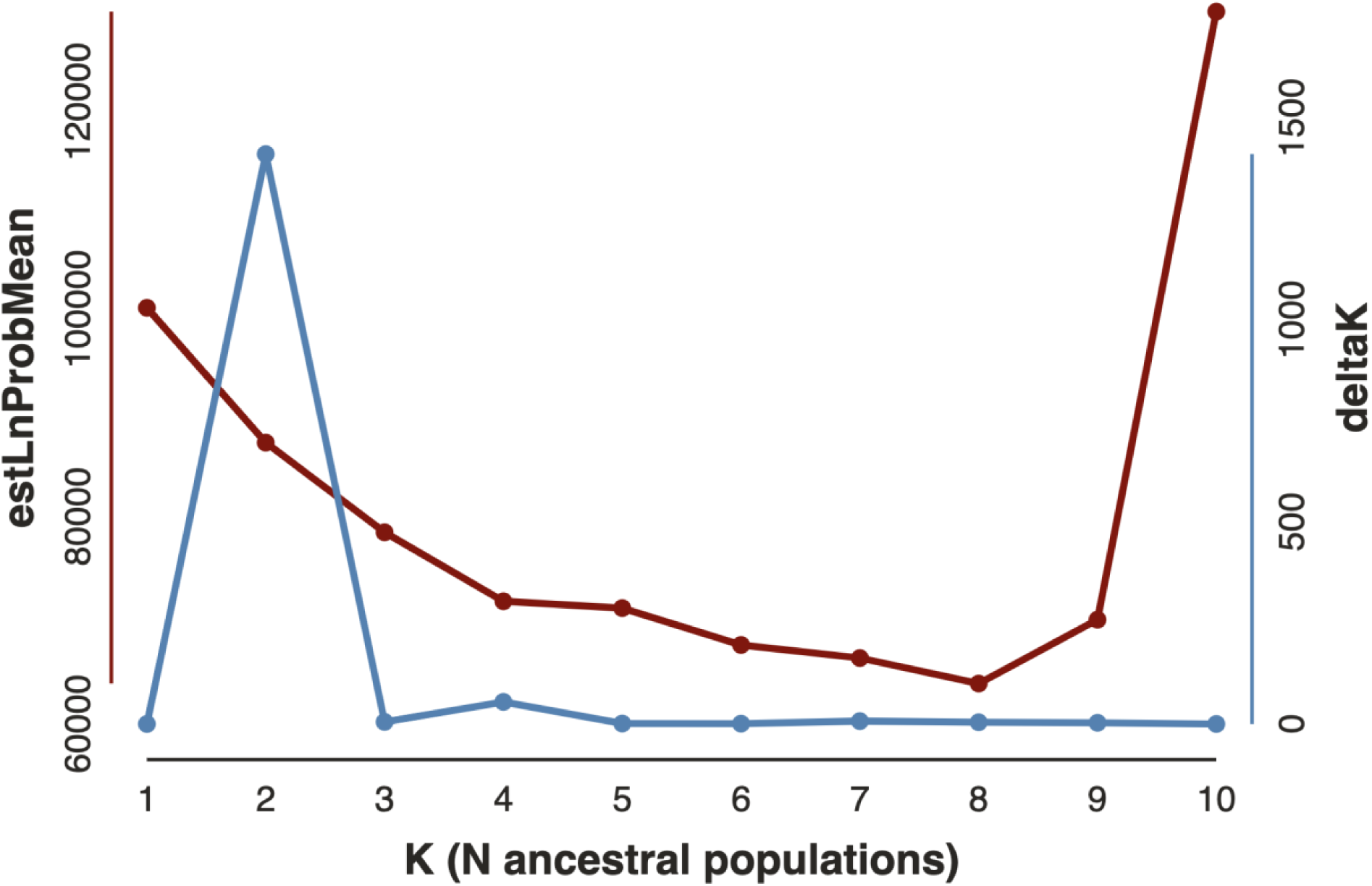
Results of two methods to select the best K value for STRUCTURE plots. The delta K method shows K=2 to be the best K, whereas the mean log probability is highest (least negative) at K=8.

**Figure S3.5.**
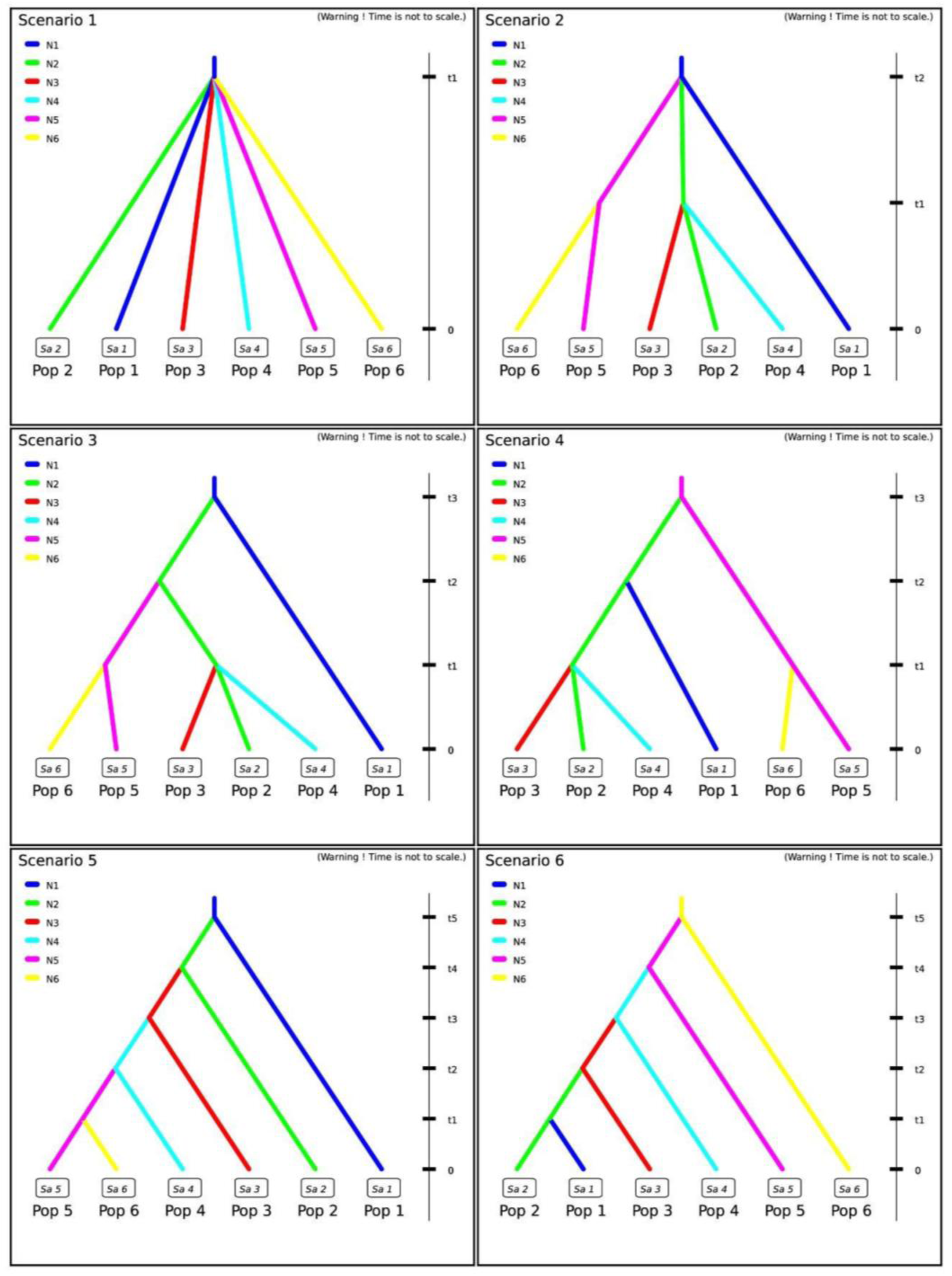
Step 1 DIYABC-RF scenarios. Pop1 = *H. liliiflorus*; Pop2 = *H. genevei*; Pop3 = *H. fragilis*; Pop4 = *H. dargentii*; Pop5 = *H. igneus*; Pop6 = *H. boryanus*.

**Figure S3.6.**
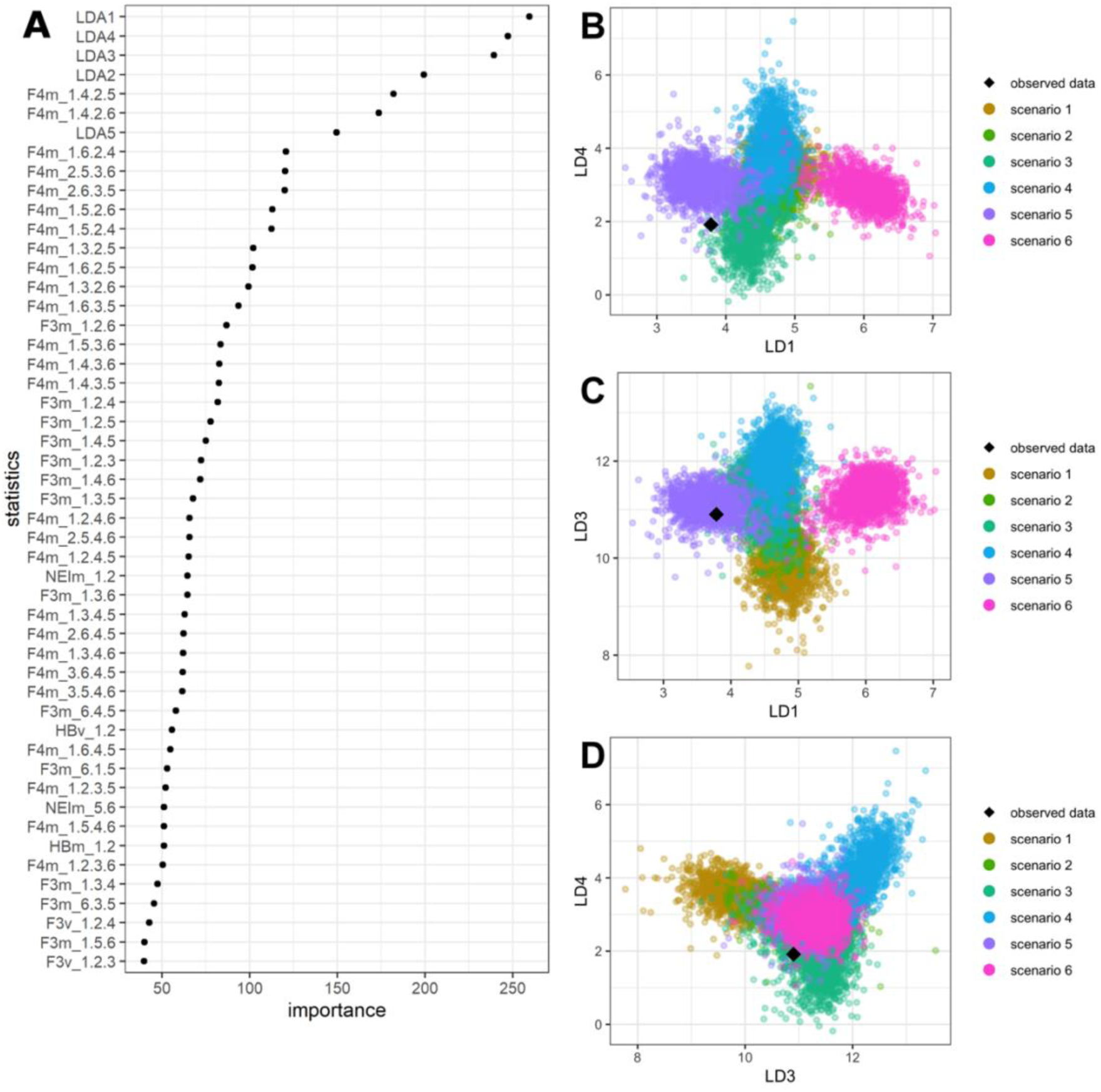
Results of DIYABC-RF linear discriminant analysis, choosing the best scenarios in step 1. A. The relative importance of each variable and linear discriminant, showing LD’s 1, 4, and 3 to be of highest importance. B-D. Plots of LD variables showing how the observed data (real dataset) was placed compared to simulated datasets at each scenario. Placement of the observed dataset in a cluster with a scenario indicates high probability that the scenario explains the observed dataset.

**Figure S3.7.**
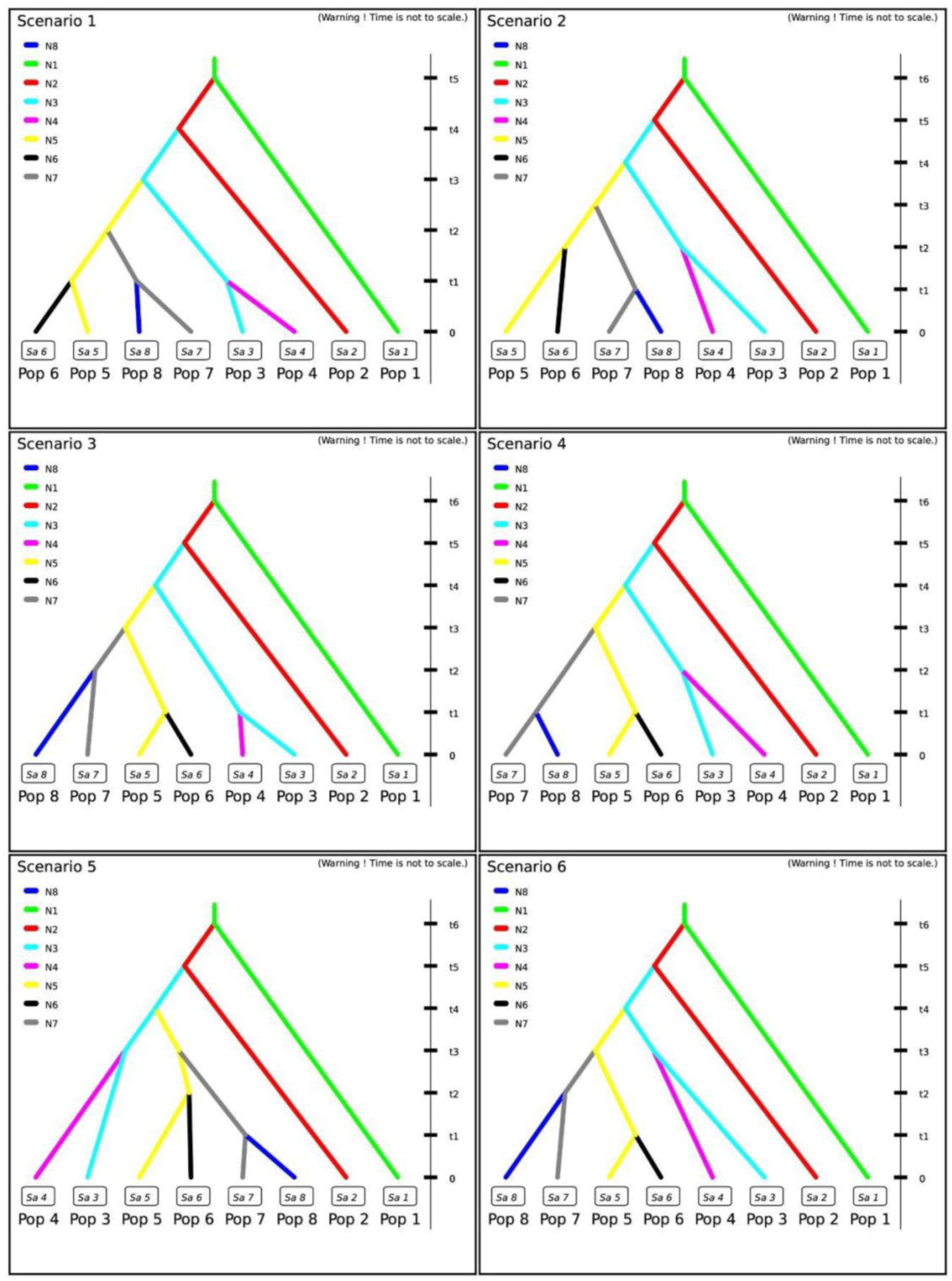
Step 2 DIYABC-RF scenarios. Pop1 = *H. liliiflorus*; Pop2 = *H. genevei*; Pop3 = *H. fragilis* North; Pop4 = *H. fragilis* South; Pop5 = *H. dargentii* North; Pop4 = *H. dargentii* South; Pop5 = *H. igneus*; Pop6 = *H. boryanus*.

**Figure S3.8.**
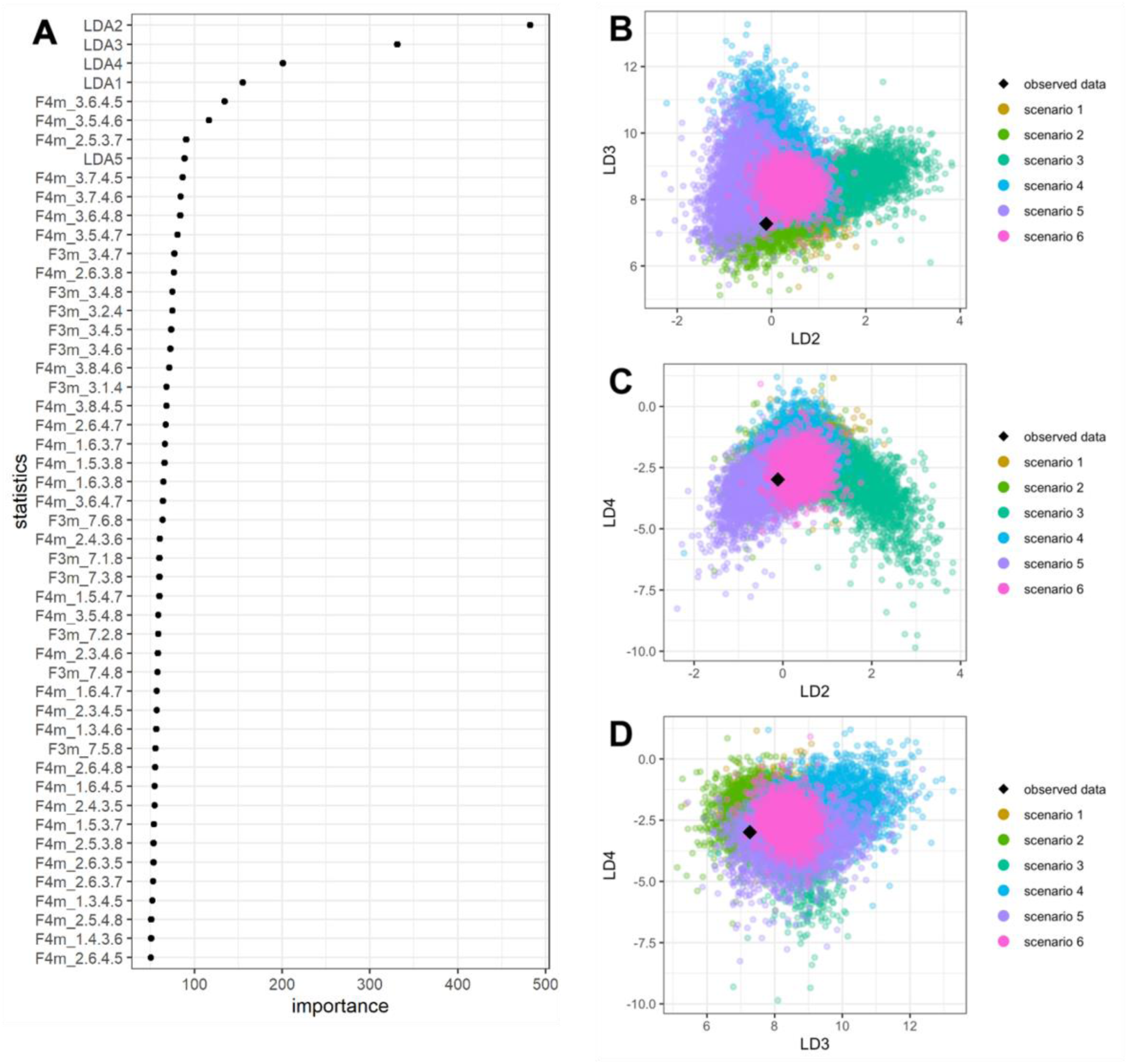
Results of DIYABC-RF linear discriminant analysis, choosing the best scenarios in step 2. A. The relative importance of each variable and linear discriminant, showing LD’s 2, 3, and 4 to be of highest importance. B-D. Plots of LD variables showing how the observed data (real dataset) was placed compared to simulated datasets at each scenario. Placement of the observed dataset in a cluster with a scenario indicates high probability that the scenario explains the observed dataset.

**Table S3.1.** Pairwise F_ST_ calculations between populations of the six Mascarene *Hibiscus* sect. *Lilibiscus* species. The northern and southern populations of *H. dargentii* and *H. fragilis* are the only populations of each species and together comprise all sampled individual. The northern and southern populations of *H. boryanus* and *H. igneus* are selected samples from the two ends of each species’ range. The bottom left of the table shows Weir and Cockerham’s F_ST_ with average values for each species at the bottom. The top right shows Nei’s F_ST_ with average values for each species at the right. Cells are colored based on variation from the mean of all F_ST_ values (0.427), with values above the mean increasingly darker red, and values below the mean increasingly darker blue.

|  | *H. boryanus* N | *H. boryanus* S | *H. dargentii* N | *H. dargentii* S | *H. fragilis* N | *H. fragilis* S | *H. genevei* | *H. igneus* N | *H. igneus* S | *H. liliiflorus* | Nei Average |
| --- | --- | --- | --- | --- | --- | --- | --- | --- | --- | --- | --- |
| *H. boryanus* N |  | 0.162 | 0.425 | 0.490 | 0.530 | 0.531 | 0.397 | 0.430 | 0.436 | 0.528 | 0.436 |
| *H. boryanus* S | 0.161 |  | 0.479 | 0.529 | 0.578 | 0.572 | 0.437 | 0.468 | 0.477 | 0.548 | 0.472 |
| *H. dargentii* N | 0.428 | 0.478 |  | 0.387 | 0.481 | 0.466 | 0.343 | 0.401 | 0.414 | 0.498 | 0.433 |
| *H. dargentii* S | 0.491 | 0.527 | 0.389 |  | 0.478 | 0.443 | 0.313 | 0.390 | 0.398 | 0.501 | 0.436 |
| *H. fragilis* N | 0.534 | 0.587 | 0.479 | 0.476 |  | 0.472 | 0.326 | 0.388 | 0.396 | 0.512 | 0.462 |
| *H. fragilis* S | 0.535 | 0.579 | 0.465 | 0.441 | 0.476 |  | 0.334 | 0.411 | 0.420 | 0.511 | 0.462 |
| *H. genevei* | 0.405 | 0.434 | 0.347 | 0.319 | 0.310 | 0.322 |  | 0.273 | 0.283 | 0.399 | 0.345 |
| *H. igneus* N | 0.450 | 0.489 | 0.415 | 0.402 | 0.392 | 0.417 | 0.265 |  | 0.077 | 0.470 | 0.367 |
| *H. igneus* S | 0.454 | 0.499 | 0.422 | 0.406 | 0.404 | 0.426 | 0.267 | 0.077 |  | 0.485 | 0.376 |
| *H. liliiflorus* | 0.511 | 0.522 | 0.475 | 0.481 | 0.467 | 0.472 | 0.360 | 0.424 | 0.432 |  | 0.461 |
| WC Average | 0.441 | 0.475 | 0.433 | 0.437 | 0.458 | 0.459 | 0.337 | 0.370 | 0.376 | 0.495 |  |

